# Profiling of transcribed *cis*-regulatory elements in single cells

**DOI:** 10.1101/2021.04.04.438388

**Authors:** Jonathan Moody, Tsukasa Kouno, Akari Suzuki, Youtaro Shibayama, Chikashi Terao, Jen-Chien Chang, Fernando López-Redondo, Chi Wai Yip, Jessica Severin, Hiroyuki Suetsugu, Yoshinari Ando, Kazuhiko Yamamoto, Piero Carninci, Jay W. Shin, Chung-Chau Hon

## Abstract

Profiling of *cis*-regulatory elements (CREs, mostly promoters and enhancers) in single cells allows the interrogation of the cell-type and cell-state-specific contexts of gene regulation and genetic predisposition to diseases. Here we demonstrate single-cell RNA-5′end-sequencing (sc-end5-seq) methods can detect transcribed CREs (tCREs), enabling simultaneous quantification of gene expression and enhancer activities in a single assay at no extra cost. We showed enhancer RNAs can be detected using sc-end5-seq methods with either random or oligo(dT) priming. To analyze tCREs in single cells, we developed *SCAFE* (Single Cell Analysis of Five-prime Ends) to identify genuine tCREs and analyze their activities (https://github.com/chung-lab/scafe). As compared to accessible CRE (aCRE, based on chromatin accessibility), tCREs are more accurate in predicting CRE interactions by co-activity, more sensitive in detecting shifts in alternative promoter usage and more enriched in diseases heritability. Our results highlight additional dimensions within sc-end5-seq data which can be used for interrogating gene regulation and disease heritability.

## Introduction

Expression of genes specifying cell-identity (i.e., cell-types and -states) is primarily controlled by the activities of their cognate CREs, mostly promoters^1^ and enhancers^2^. CREs are highly enriched in disease-associated variants^3^, reflecting the importance of gene regulation in diseases. Therefore, understanding the cell-identity-specific CRE activities not only helps to decipher the principles of gene regulation^4,5^, but also the cellular contexts of genetic predisposition to diseases^6^. While gene expression can be quantified with single-cell RNA-sequencing methods (sc-RNA-seq)^7,8^, profiling of CREs primarily relies on single-cell Assay for Transposase Accessible Chromatin using sequencing (sc-ATAC-seq)^9,10^, which measures the accessibility of chromatin regions in a binary manner (i.e., accessible or non-accessible)^11^. Several methods were developed for joint profiling of gene expression and chromatin accessibility within the same cell^12–15^, allowing the prediction of CRE interactions with their target genes (i.e., enhancer-to-promoter (EP) interactions), through cell-to-cell co-variations of their activities^13,16^. However, the close-to-binary nature and excess sparsity of chromatin accessibility data render the analyses of individual CREs in single cells challenging^17^. Also, a substantial fraction of accessible CREs that are distant from annotated promoters (i.e., distal aCREs) do not show the epigenomic features of active enhancers^18^. While an unknown fraction of non-enhancer distal aCREs could be regulatory, e.g., insulators^19^ or silencers^20^, their overall relevance in gene regulation remains elusive.

Alternatively, measuring the transcription at CRE can be used as a proxy for their activity^2^, which can be achieved by sequencing the 5′ends of RNA^21^ representing the transcription start sites (TSS) within CREs^1^. Such measurement is highly quantitative and is ranked as the top feature for predicting active EP interactions in a machine-learning approach, compared to other epigenomic features^22^. In fact, the co-variation of transcription signals between CREs were shown to accurately predict individual cell-type-specific EP interactions^23^. In addition, endogenous transcription at a distal CRE is highly correlated with its ability to function as an enhancer in transgenic enhancer assays^24^. These observations suggest distal CRE identified and quantified by transcription evidence, compared to solely chromatin accessibility evidence, could be more relevant to enhancer activation of gene expression.

Previously, we demonstrated the application of sc-end5-seq in the integrated fluidic circuit-based C1 platform (Fluidigm) for detection of known TSS in hundreds of single cells^25^. In this study, we evaluated the sc-end5-seq methods on the droplet-based Chromium platform (10x Genomics), with random and oligo(dT) priming, for *de novo* discovery and quantification of tCREs in thousands of single cells. Unexpectedly, both random and oligo(dT) priming methods detected enhancer RNAs, which are generally thought to be non-polyadenylated (non-polyA)^2^. A major challenge in *de novo* discovery of tCREs from sc-end5-seq data is artifactual TSS^26,27^ from template switching (TS) reactions. Therefore, we have devised a multiple logistic regression classifier to effectively minimize artifactual TSS. Applying both sc-end5-seq and sc-ATAC-seq to peripheral blood mononuclear cells (PBMC) under stimulation, we compared the performance of tCREs and aCREs in 1) identification of cell-type-specific CREs, 2) detection of stimulation-induced transcription factor (TF) activities, 3) detection of shifts in alternative promoter usage, 4) prediction of CRE interactions by co-activity, 5) enrichment in diseases heritability and 6) functional interpretations of disease-associated variants. Finally, we developed *SCAFE*, a command-line tool to define genuine tCREs and predict their interactions from RNA-5′end-sequencing data.

## Results

### Assessing the performance of 3′end and 5′end sc-RNA-seq methods

While the sc-end5-seq method on the Chromium platform (10x Genomics) is primed with oligo(dT) (sc-end5-dT), we modified the protocol with random hexamer priming (sc-end5-rand), aiming for better detection of non-polyA RNAs (Methods)^28,29^. We then performed both sc-end5-dT and sc-end5-rand methods, along with the oligo(dT) primed 3′end sc-RNA-seq method (sc-end3-dT), on human dermal fibroblasts (DMFB) and induced pluripotent stem cells (iPSC) (Supplementary Fig. 1a). For comparison, bulk-CAGE, bulk-RNA-seq, and bulk-ATAC-seq were also performed on both cell lines (Fig. 1). In the following section, we focus on iPSC for the sake of brevity (Fig. 2; Supplementary Fig. 2 for DMFB).

**Fig. 1.**
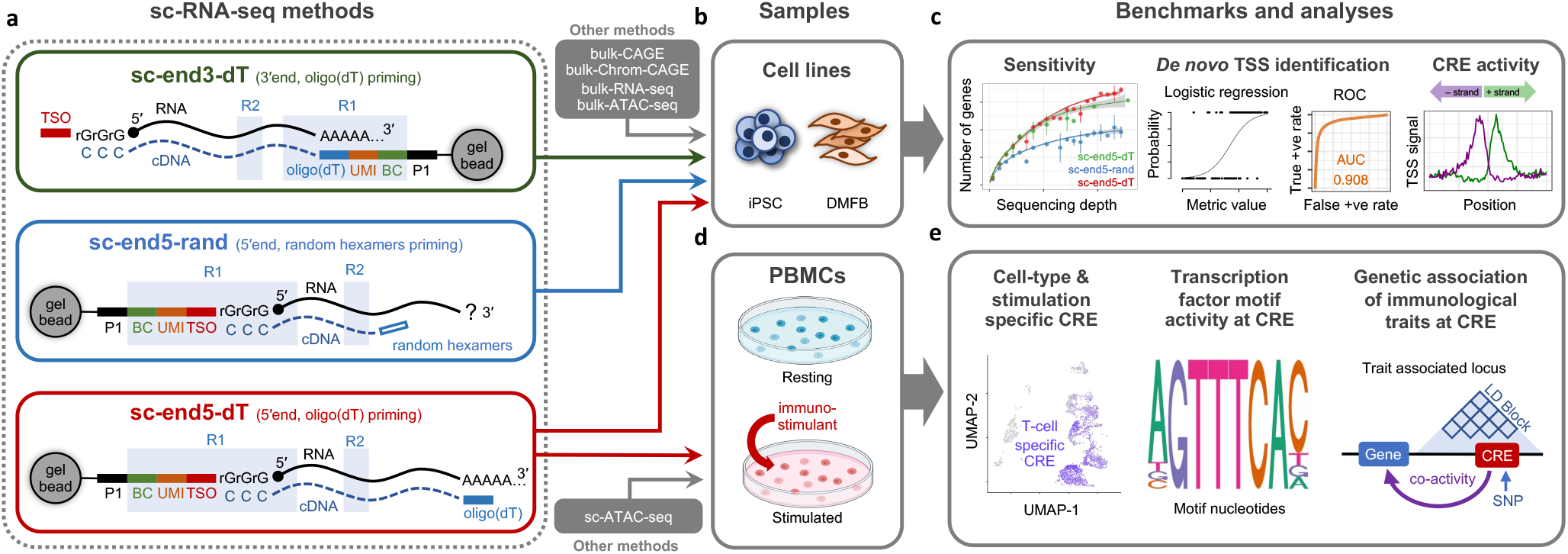
Overview of the experimental designs and benchmark analysis. **a**, sc-RNA-seq methods used in this study. sc-end5-rand is a custom method, and the other two methods are original methods on Chromium platform (10x Genomics). *BC*: cell barcode, *UMI*: unique molecular identifier, *TSO*: template switching oligonucleotide, *R1*: read 1, *R2*: read 2, *P1*: sequencing primer 1. **b**, Two cell lines, iPSC and DMFB, were used to compare the performance of the three sc-RNA-seq methods, with matched bulk transcriptome and epigenome datasets. **c**, The datasets from **(b)** were used for sensitivity assessment, *de novo* TSS identification, CRE activity detection. **d**, PMBCs, at resting and stimulated states, were profiled using sc-ATAC-seq and sc-end5-dT. **e**, The datasets from **(d)** were compared in terms detection of cell-type/stimulation specific CRE, transcription factor motif activity and genetic association of traits.

**Fig. 2.**
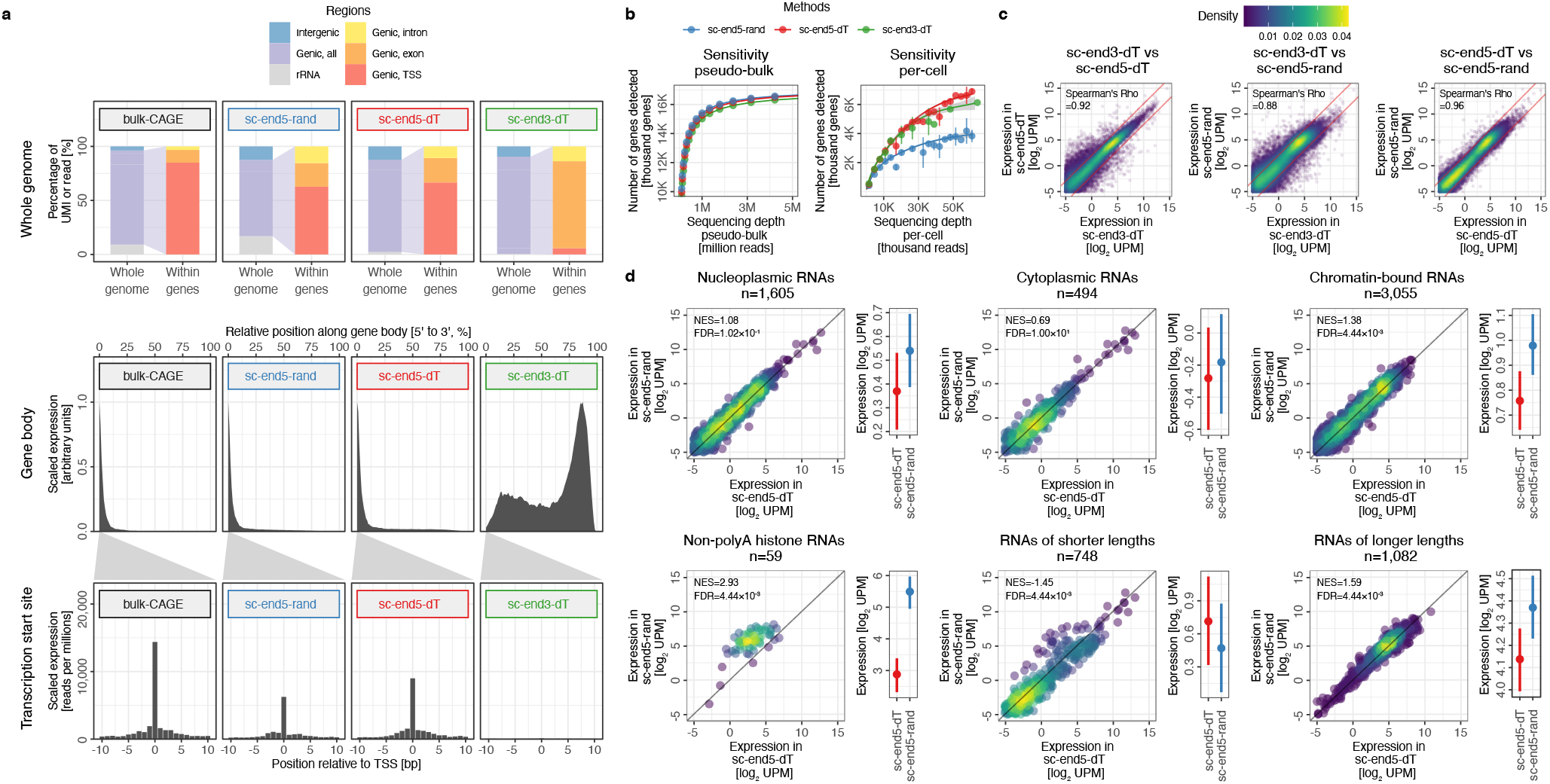
Performance of sc-RNA-seq methods. **a**, Distribution of reads from bulk-CAGE and sc-RNA-seq methods. *Top*, distribution of reads in the whole genome; *Middle*, distribution of reads along the gene body; *Bottom*, distribution of reads in surrounding annotated TSS. **b**, Sensitivity of gene detection in pseudo-bulk (*left*) and in single cells (*right*) across sequencing depth. Error bars represent standard deviation. The genes that are detected in bulk-RNA-seq were used to define the scope. **c**, Correlation of gene expression levels between the pseudo-bulk data of the three sc-RNA-seq methods. *Red line*, ±2-fold differences. *UPM*, UMI per millions. Color represents the density of points. **d**, Differences in the expression levels of RNAs with various properties between sc-end5-rand and sc-end5-dT. Gene Set Enrichment Analysis (GSEA) was performed on each RNA set; NES and FDR, normalized enrichment score and false discovery rate of GSEA. Color represents the density of points. A positive NES value with FDR <0.05 refers to a significantly higher abundance of an RNA set in sc-end5-rand. *Right*, mean and standard errors.

First, we assessed the global distributions of reads. As expected, in sc-end5-rand more reads are mapped to ribosomal RNA (rRNA) (∼15%) than in sc-end5-dT (∼2%) (Fig. 2a, *upper panel, whole genome*). When considering genic reads, the percentage of reads mapped to TSS is lower in both sc-end5-seq (∼60%) than in bulk-CAGE (∼85%) (Fig. 2a, *upper panel, within gene*), reflecting the greater extent of artefacts in sc-end5-seq, as discussed in the next section. We also noted the genic reads of both sc-end5-seq methods and bulk-CAGE were strongly enriched at the 5′end of genes (Fig. 2a, *middle panel*) and peaked precisely at the annotated TSS (Fig. 2a, *lower panel*), suggesting both sc-end5-seq methods can precisely pinpoint TSS.

Next, we assessed gene detection sensitivity. In pooled single cells (i.e., pseudo-bulk), all three sc-RNA-seq methods showed similar sensitivities (Fig. 2b, *left panel*). However, when considered per-cell, both oligo(dT)-primed methods (i.e., sc-end3-dT and sc-end5-dT) detected ∼30% more genes than the random-primed method (i.e., sc-end5-rand) at matched sequencing depths (Fig. 2b, *right panel*). This might be explained by lower complexity of the sc-end5-rand per-cell libraries, attributed to its higher rRNA read percentage and higher reads per unique molecular identifier (UMI) (Supplementary Fig. 1). Overall, the pseudo-bulk expression level of genes among the three sc-RNA-seq methods are highly correlated (Fig. 2c), allowing datasets from these three sc-RNA-seq methods to be robustly integrated (Supplementary Fig. 3), opening the possibility of joint-analyses of sc-end5-seq datasets with ample sc-end3-dT public datasets.

To further examine the differences between the two priming methods, we tested for the enrichment of subcellular compartment-specific RNAs (Methods), non-polyA histone RNAs^30^, and long or short RNAs (Methods). In the genes with higher abundance in sc-end5-rand compared to sc-end5-dT, we observed strong enrichment of non-polyA histone RNAs (FDR <0.005, Fig. 2d). This is supported by the enrichment of chromatin-bound RNAs (FDR <0.005, Fig. 2d), which include many nascent RNAs and non-polyA RNAs^31^. The significant enrichment of long RNAs (FDR <0.005, Fig. 2d) might be attributed to the higher reverse transcription efficiency of random priming within the body of the longer transcripts, in contrast to oligo(dT) priming which mainly from transcript 3′ends. Unexpectedly, sc-end5-dT also detected non-polyA histone RNAs with default expression (Fig. 2d), suggesting potential internal priming at A-rich sequences in sc-end5-dT, which has been also observed extensively in sc-end3-dT method^32,33^. In summary, these observations suggest a comparable performance in gene detection for the three sc-RNA-seq methods, with sc-end5-rand showing slightly lower per-cell sensitivity and sc-end5-dT showing unexpected detection of non-polyA RNAs.

### TSS identification using sc-end5-seq methods

Previous reports suggested a fraction of TSS identified based on read 5′ends from TS reactions may not be genuine^26,27^, attributed to artefacts including strand invasion^27^ and other sources (e.g., sequence biases)^34^. This results in excessive artifactual TSS, especially along the gene body, known as “exon painting”^35^. While a fraction of “exon painting” reads could be attributed to cleavage and recapping^36^, their exact molecular origins remain elusive. Here we developed a workflow in *SCAFE* to identify genuine TSS (Supplementary Fig. 4).

First, we filtered strand invasion artefacts based on the complementarity to TS oligo sequence^34^ and found more strand invasion artefacts in sc-end5-rand (∼5% reads) than in sc-end5-dT (∼3% reads) (Supplementary Fig. 5). The filtered reads were then grouped as TSS clusters. We observed a substantially higher proportion of TSS clusters along the gene body in both sc-end5-seq methods than bulk-CAGE (Supplementary Fig. 6), consistent with the fact that “exon painting” is more prevalent in TS-based methods^26^. We benchmarked the properties of TSS clusters (Fig. 3a) and devised a classifier for genuine TSS (Methods) (Fig. 3b). Here we focus on the sc-end5-dT iPSC dataset for simplicity. First, the UMI counts within the TSS cluster (cluster count) performed the worst (Area Under Receiver Operating Characteristic (ROC) Curve (AUC)=0.641) (Fig. 3a), and its performance decreased with sequencing depth (Fig. 3c). Two other common metrics, UMI count at TSS summit (summit count, AUC=0.725) and within ±75nt flanking its summit (flanking count, AUC=0.737) performed only marginally better than the cluster count (Fig. 3a,c), suggesting these commonly used metrics are at best mediocre classifiers for TSS. As “exon painting” artefacts are positively correlated with transcript abundance, making count-based thresholds poor performers, we examined other metrics that are independent of RNA expression levels, including UMI counts corrected for background expression (corrected expression, Methods) and percentage of reads with 5′mismatched G^26^ (unencoded-G percentage, Methods). Notably, we found both metrics performed well across sequencing depths with AUC >0.9 (Fig. 3c).

**Fig. 3.**
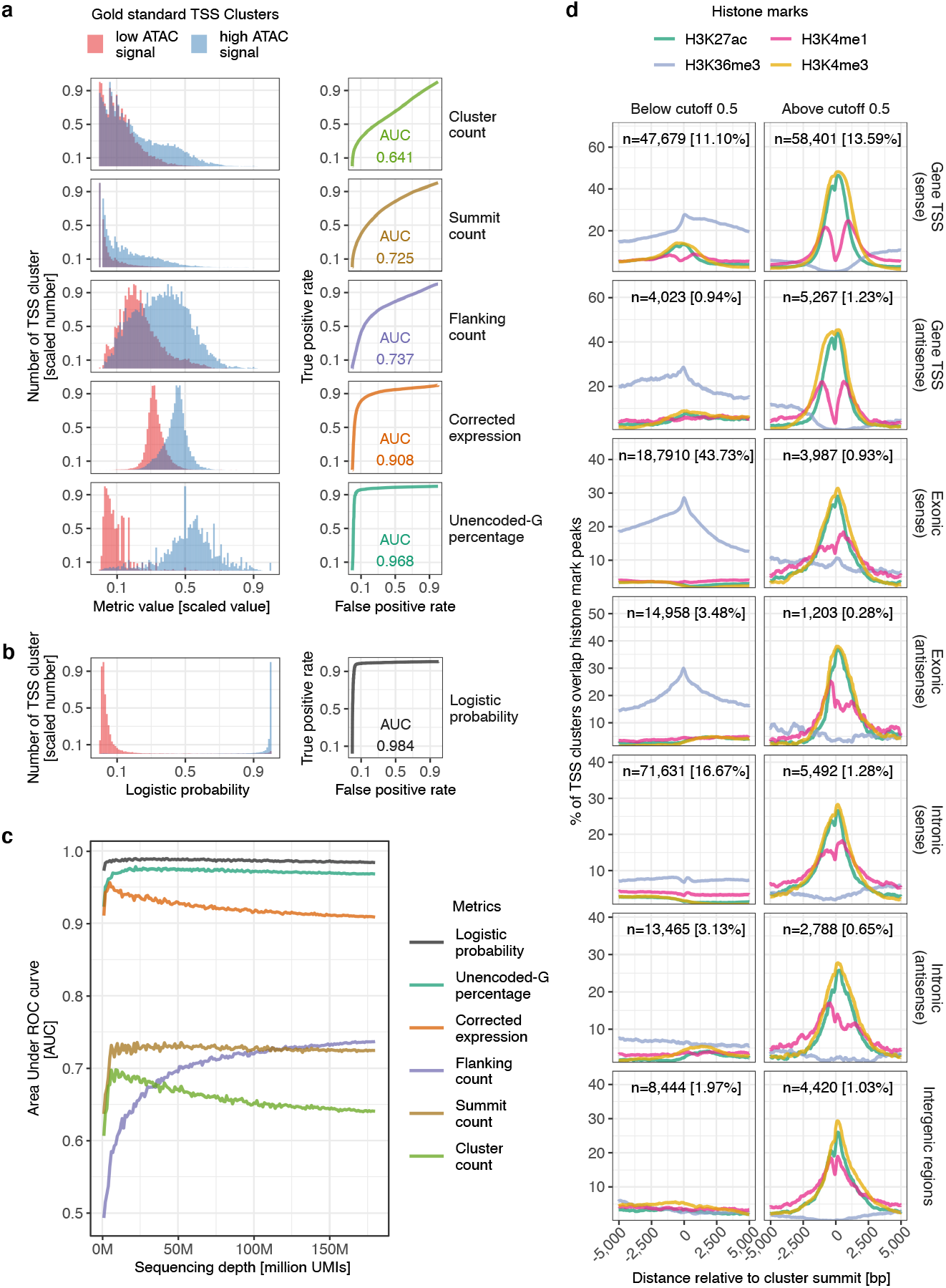
*De novo* identification of genuine TSS. **a**, Properties of gold standard TSS clusters (*left*) and their performance as a TSS classifier measured as Area Under Receiver ROC Curve (AUC) (*right*). **b**, Logistic probability of gold standard TSS clusters (*left*) and its performance as a TSS classifier measured as AUC (*right*). **c**, Performance of various metrics as a TSS classifier in (**a**) and (**b**) across various sequencing depth. **d**, Histone marks at TSS clusters with logistic probability below (*left*) or above (*right*) 0.5 cutoff, at annotated gene TSS, exonic or intronic regions in sense or antisense orientations, or otherwise intergenic regions. n, number of TSS clusters. %, percentage of TSS clusters in all genomic locations regardless of logistic probability thresholds.

We then used multiple logistic regression to devise a TSS classifier. We found the combination of flanking count, unencoded-G percentage and corrected expression is paramount to achieve the best performance, with AUC >0.98 across sequencing depths (Fig. 3b,c). Its accuracy is high and robust for TSS clusters located in various genomic regions and across a wide range of cutoffs (Supplementary Fig. 7a), which is well-validated by chromatin accessibility, promoter motifs, CpG island, sequence conservation (Supplementary Fig. 7b-f) and histone marks (Fig. 3d). At the default cutoff of 0.5, ∼98% of sense exonic TSS clusters were removed (Fig. 3d, *3rd row*). These removed TSS clusters are void of marks for active CREs (e.g., H3K27ac, H3K4me1 and H3K4me3) but overlap marks for transcription elongation (e.g., H3K36me3), suggesting our TSS classifier effectively removed “exon painting” artifacts. In addition, the TSS clusters located at gene TSS are marked with a bimodal H3K4me1 pattern which indicates active promoters, in contrast to the others that are marked with relatively unimodal H3K4me1 pattern which indicates active enhancers^37,38^. In summary, our TSS classifier robustly distinguishes genuine TSS from artifacts. These data can be browsed online under “TSS in Single Cells” and “Benchmarking TSS Clusters” themes (Data availability).

### Defining tCRE using sc-end5-seq methods

tCREs were defined by merging closely located TSS clusters and classified as either proximal or distal based on their distance to annotated gene TSS (Fig. 4a). ‘Proximal’ tCRE can be interpreted as promoters of genes and their upstream antisense transcripts (PROMPTs)^39^. ‘Distal’ tCRE can be interpreted as mostly enhancers^40^, with an unknown, but likely minor, fraction of them as unannotated promoters (e.g., alternative promoters). To benchmark the sensitivity of tCRE detection, we performed bulk-CAGE on chromatin-bound RNA (bulk-Chrom-CAGE), which captures the 5′ends of nascent transcripts for sensitive detection of short-lived RNAs (e.g., enhancer RNAs)^31^ and can be viewed as a permissive baseline for their detection. First, we found similar proportions of distal tCREs in sc-end5-dT (∼10%) and sc-end5-rand (∼12%) (Fig. 4b, in *all tCRE*), suggesting a similar sensitivity of enhancer RNA detection in both methods. Amongst all distal tCREs, the proportions of exonic, intergenic and intronic were similar between the bulk and single-cell methods (Fig. 4b, in *distal tCRE*). Considering the excessive exonic TSS cluster in sc-end5-seq before filtering (Supplementary Fig. 6), the TSS classifier appeared to effectively minimize “exon painting” artefacts.

**Fig. 4.**
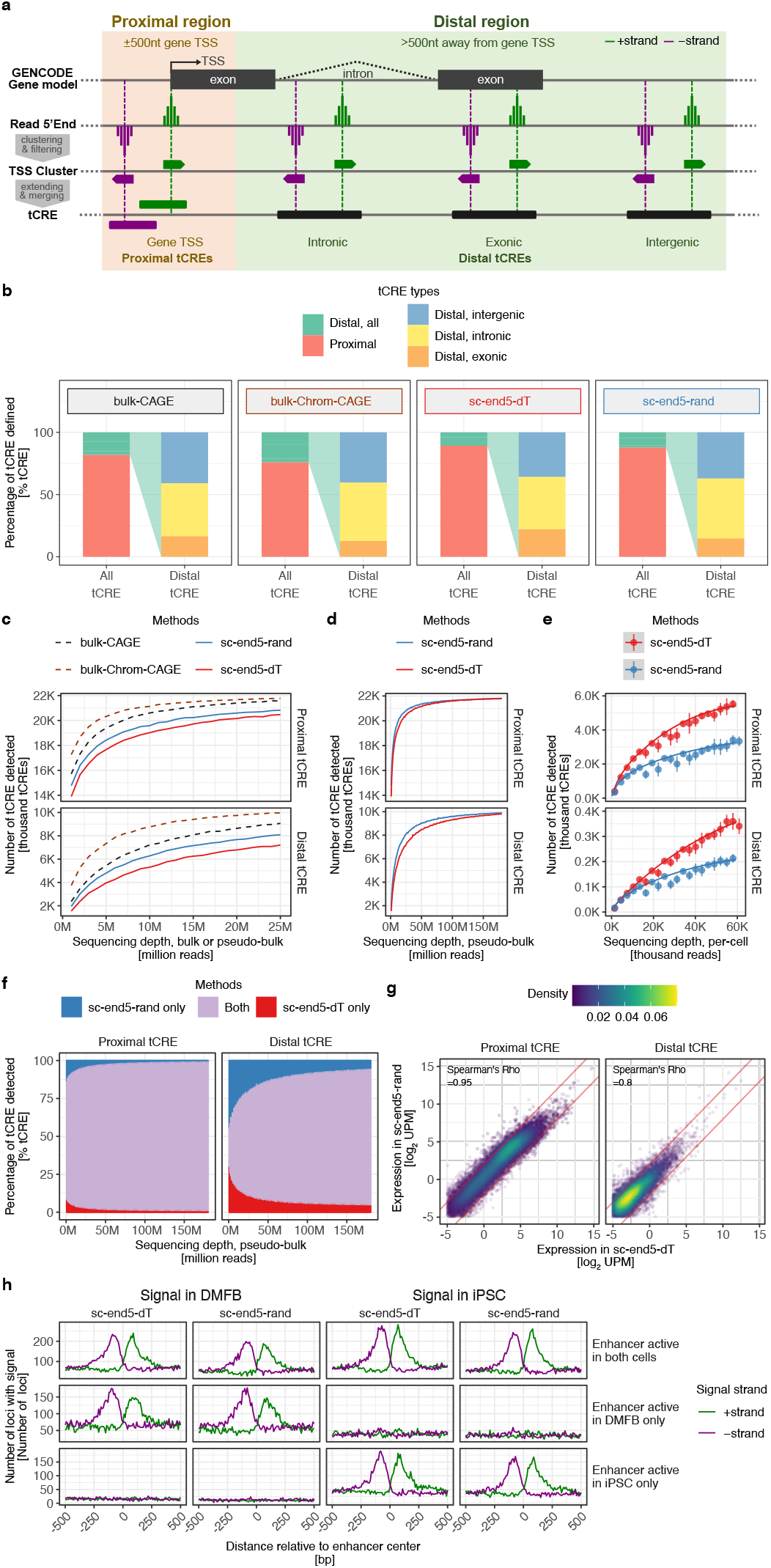
Definition and properties of tCRE. **a**, Defining tCRE by merging closely located TSS clusters. Distance to gene TSS was used as the criteria to define proximal or distal TSS clusters and tCREs. Proximal and distal TSS clusters were merged in stranded and strandless manner, respectively. Distal tCREs are further classified as intronic, exonic, or otherwise intergenic. **b**, Proportion of tCREs types defined from sc-end5-dT and sc-end5-rand pseudo-bulk, compared to bulk-CAGE and bulk-Chrom-CAGE. All four libraries were subsampled to 25 million reads. **c**, Sensitivity of tCRE detection in sc-end5-dT and sc-end5-rand pseudo-bulk, compared to bulk-CAGE and bulk-Chrom-CAGE, from 1 to 25 million reads. **d**, Sensitivity of tCRE detection in sc-end5-dT and sc-end5-rand pseudo-bulk, from 1 to 150 million reads. **e**, Sensitivity of tCRE detection in sc-end5-dT and sc-end5-rand in single cells, from 1,000 to 60,000 reads per cell. Error bars represent standard deviation. **f**, Proportion of overlap in tCRE detected in sc-end5-seq pseudo-bulk from 1 to 150 million reads. **g**, Correlation of tCRE levels between the pseudo-bulk data of the two sc-end5-seq methods. *Red line*, ±2-fold differences. *UPM*, UMI per millions. **h**, TSS signal of sc-end5-dT and sc-end5-rand at bidirectionally transcribed enhancer loci defined in bulk-CAGE in iPSC and DMFB.

Assessing the sensitivity of tCRE detection, we found bulk-Chrom-CAGE has the highest sensitivity, while both sc-end5-seq methods detected ∼50% to ∼80% of those detected by bulk methods at matched sequencing depths (Fig. 4c). In pseudo-bulk, sc-end5-rand seemed slightly more sensitive at lower depths (Fig. 4d, at ∼50M), but both methods performed similar at higher depths (Fig. 4d, at ∼150M). When considering per-cell, however, sc-end5-dT is substantially more sensitive than sc-end5-rand (Fig. 4e). This might be explained by lower complexity in sc-end5-rand libraries as discussed earlier in Fig. 2b. Unexpectedly, the distal tCREs identified in both methods are largely overlapping (Fig. 4f) and their expression levels are highly correlated (Fig. 4g), considering a substantial fraction of distal tCREs are enhancer RNAs that lack polyA tails^2^. To further investigate this, we examined the bidirectionality of transcribed enhancer loci in DMFB and iPSC (defined by bulk-CAGE as previously described^2^). Both sc-end5-dT and sc-end5-rand recapitulated the bulk-CAGE defined bidirectional transcription pattern in a cell-type-specific manner (Fig. 4h), supporting that both sc-end5-seq methods detected enhancer RNAs with similar sensitivity. The unexpected detection of enhancer RNAs by sc-end5-dT could be attributed to the potential internal priming^32,33^. Taken together, in view of their similar pseudo-bulk performances (Fig. 4d,f-h) and the superior per-cell performance of sc-end5-dT (Fig. 4e), we performed sc-end5-dT and sc-ATAC-seq in PBMC for the comparison of tCRE and aCRE.

### Comparing tCRE and aCRE in PBMC

We next defined tCREs (n=30,180) and aCREs (n=157,055) in PBMCs in stimulated or resting cell-state (Methods) (Fig. 1; Supplementary Fig. 1). Gene-based cell-type annotations were transferred from the tCRE cells to aCRE cells using CCA^41^ (Supplementary Fig. 8). UMAPs based on tCRE or aCRE show similar clustering of cell-types and excellent integration of cell-states (Fig. 5a). Examining a subset of aCREs with cell-type-specific chromatin accessibility (Methods, Fig. 5b, *top row*), we found concordant patterns of cell-type-specific RNA transcription at the overlapping tCREs (Fig.5b, *bottom row*). To examine TF activities, we applied *ChromVAR*^42^ to both aCRE and tCRE to define cell-type-specific TF motifs (Methods). The cell-type-specific TF motifs defined based on aCRE and tCRE are significantly concordant in most cell-types (Supplementary Fig. 9a, Fisher’s exact test, P <0.05). Clustering of cell-types using TF motif activities appears to be consistent within broad categories with co-clustering of monocytes, lymphocytes and cytotoxic T-cells between aCRE and tCRE (Supplementary Fig. 9b). We further examined the activation of TF upon stimulation (Methods) and observed a generally consistent upregulation of TF activities between aCREs and tCREs (Fig. 5c, mean Pearson’s r=0.84), mostly driven by *JUN/FOS* related motifs that are components of the early immune responses. These results suggest both tCRE and aCRE can recover cell-type and cell-state-specific features of gene regulation (e.g., CRE and TF activities). These data can be browsed online under “Cell-Type-Specific CREs” (Data availability).

**Fig. 5.**
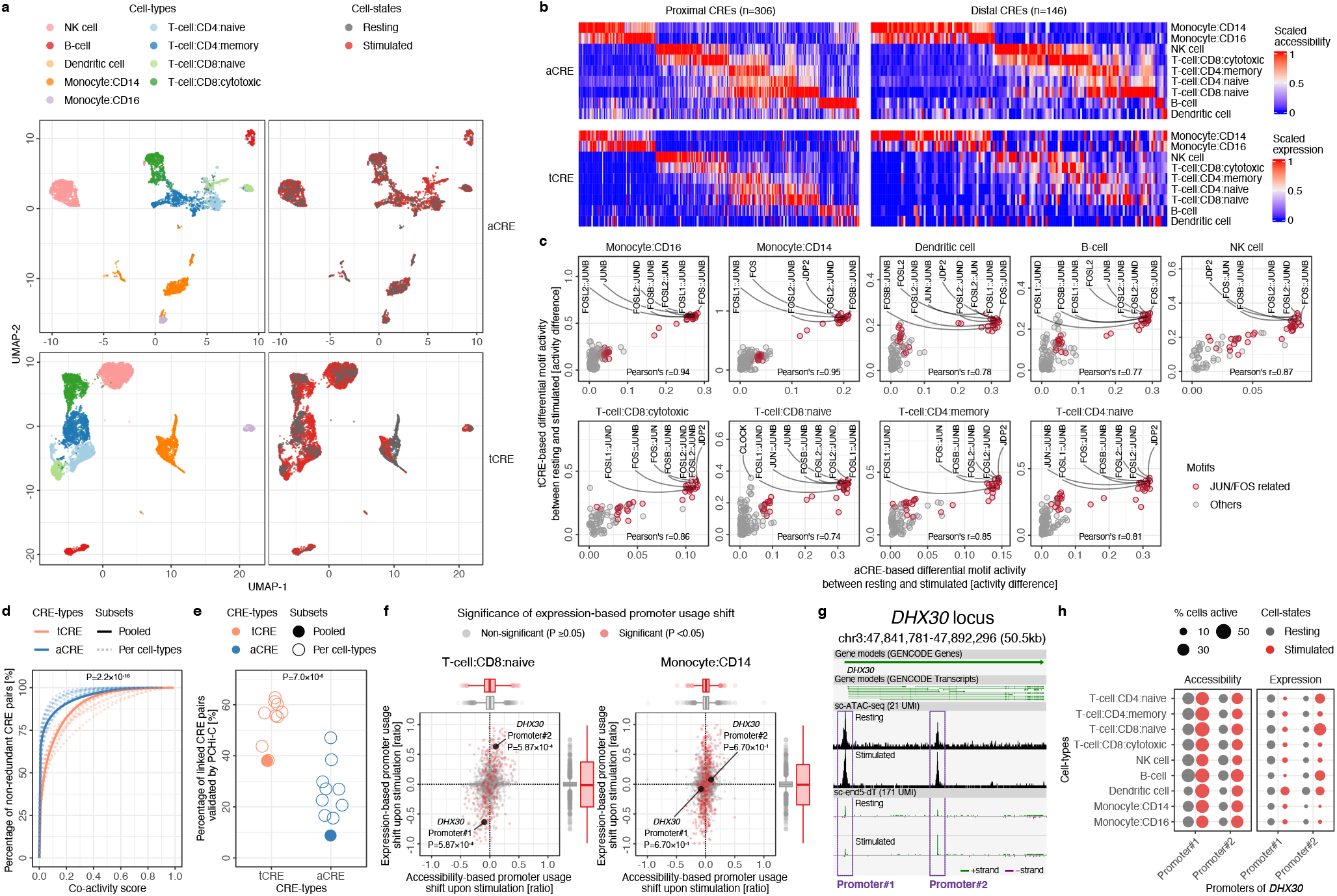
Comparison of tCRE and aCRE in PBMCs. **a**, UMAP of cells based on aCREs *(upper*) and tCREs (*lower*), colored by cell-types (*left*) or cell-states (*right*). **b**, Heatmap of cell-type specific CREs. Only CRE loci overlapped with both aCREs and tCREs were selected. Column orders were sorted based on cell-type specificity measured in aCRE. *Upper*, aCRE; *Scaled accessibility*, ratio of cells with open aCRE, normalized to the maximum value per loci. *Lower*, tCRE; *Scaled expression*, mean expression per cell-type normalized to the maximum value per loci. **c**, Motif activity difference between resting and stimulated cells in aCRE (*x-axis*) and tCRE (*y-axis*) per cell-type. *JUN/ FOS* family motifs are highlighted. **d**, Distribution of co-activity scores for tCRE (*orange*) and aCRE (*blue*) within each cell-type (*dashed lines*) and all cells pooled (*solid line*). Kolmogorov-Smirnov test statistic (P) for difference of distribution in all cells pooled shown. **e**, Percentage of linked CRE pairs (co-activity score ≥0.2) validated (by PCHi-C) for tCRE (*orange*) and aCRE (*blue*), for per cell-type (*hollow circles*) and for all cells pooled (*solid circles*). T-test for difference of tCRE and aCRE means shown. **f**, Shifts in alternative promoter usage upon stimulation for genes with multiple proximal tCRE. *X-axis*, change in accessibility (ratio of proportion of signal in sc-ATAC-seq) within tCRE upon stimulation; *Y-axis*, mean change in expression (ratio of proportion of signal in sc-end5-dT) of tCRE across metacells (k=50) upon stimulation. P, *t-*test for change in tCRE usage between metacells shown. **g**, Alternative promoter usage shift at *DHX30* locus, modified from Zenbu genome browser view (Data availability). **h**, Cell-type specific shift in alternative promoter usage at *DHX30* locus. Proportion of cells with accessible aCRE (*left*) and transcribing tCRE (*right*) colored by stimulation state.

Co-activity of a pair of CREs can be used to predict their physical interactions^16^. Here we compared the accuracy of tCREs and aCREs in prediction of interacting CREs, using the co-activity estimated in *Cicero*^16^, benchmarked against promoter-capture Hi-C (PCHi-C)^43^ (Methods). Co-activity scores were estimated separately using cells within individual cell-types (cell-type sets) or all cells (pooled set). Here, we focus on a subset of CREs that is overlapping between tCRE and aCRE (Methods). First, we observed significantly higher co-activity scores for tCRE-pairs than aCRE-pairs (Fig. 5d, P <2.2×10^−16^, Kolmogorov-Smirnov test for the pooled set, *solid line*). At co-activity scores ≥0.2, we found the linked tCRE-pairs are significantly more likely to be validated by PCHi-C (∼40%) than the linked aCRE-pairs (∼10%) (Fig. 5e, P <7×10^−6^, paired *t*-test for the cell-type sets). Despite the lower number of tCRE in comparison to aCRE, these results suggest tCREs are more accurate in predicting CRE interactions by co-activity. These data can be browsed online under “Interactions between tCREs” (Data availability).

Alternative promoter usage is an important mechanism to increase transcriptome diversity for generation functionally distinct isoforms^44^. We found 123 genes with significant shifts in proximal tCRE usage upon stimulation in at least one cell-type (FDR <0.05 in *t*-test). We then examined the chromatin accessibility at the corresponding tCREs and observed only minimal extent of shifts in accessibility (Fig. 5f, *horizontal box plot, top*). Highlighting the *DHX30* locus (Fig. 5g), in T-cell:CD8:naïve, its expression shifted from Promoter#1 to Promoter#2 upon stimulation, whereas in Monocyte:CD14, no shift was observed (Fig. 5f,h; Supplementary Fig. 10). In contrast, the chromatin accessibility at the two promoters remains mostly constant between the two cell-states in all cell-types (Fig. 5h). These results suggest tCREs are generally more sensitive in detecting shifts in alternative promoter usage upon cell-state changes. These data can be browsed online under “Alternative Promoter Usage” (Data availability).

### Enrichment of trait-associated variants in tCRE

For interpretation of genetic predisposition, we examined the enrichment of trait heritability^45^ in CREs from PBMCs. For comparison with aCRE, we used tCRE defined with TSS clusters at default and lenient logistic probability thresholds (Methods). As expected, we found both tCREs and aCREs are enriched in hematologic and immunologic traits, but generally not in psychiatric and metabolic traits (Fig. 6a, *top row*). The pattern is similar when considering proximal and distal CREs separately (Fig. 6a, *middle and bottom row*), implying that the distal tCREs are biologically relevant. In addition, the enrichment in default tCREs is generally higher than that of lenient tCREs, particularly for distal tCREs (Fig. 6a), suggesting a higher proportion of default tCREs are biologically relevant. However, we also noticed the default tCRE are less sensitive in terms of reaching statistical significance, which can be attributed to the smaller number of SNPs in default tCRE leading to larger estimates of standard error as reported^46^. For more statistical power, we thus used lenient tCREs in the following analyses.

**Fig. 6.**
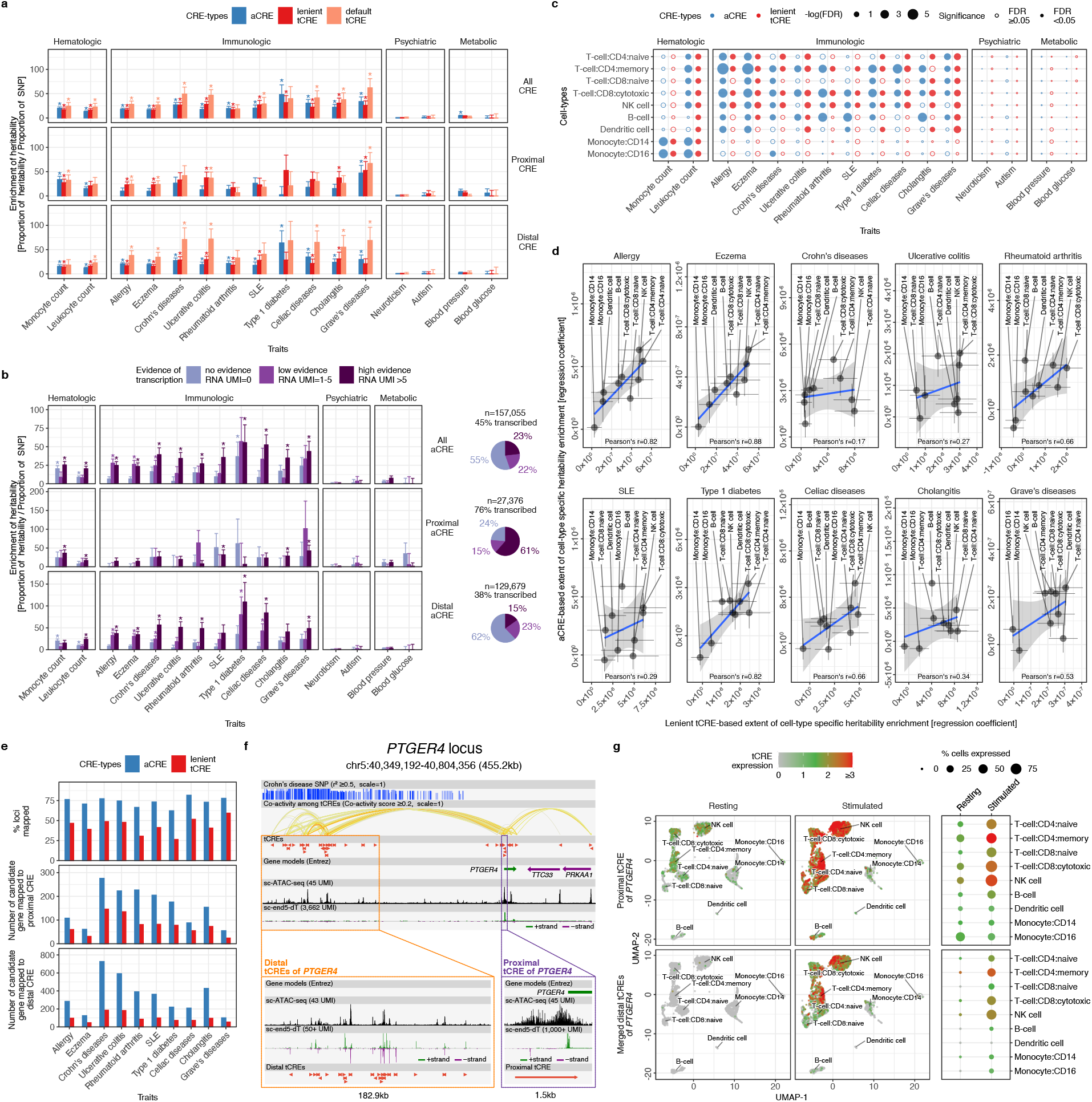
Disease-associated variants at tCRE and aCRE in PBMCs. **a**, Enrichment of heritability in various CRE-types. *Y-axis*, enrichment of heritability is measured as the ratio of proportion of heritability to proportion of SNP, estimated by LDSC. *Error bars*, standard error of the estimate. *Asterisks*, significant enrichments with FDR <0.05. **b**, Enrichment of heritability in aCREs with various levels of evidence of transcription. *Y-axis, error bars*, and *asterisks* are the same as in **(a). c**, Enrichment of heritability in cell-type specific CREs. *Solid circles*, significant enrichments with FDR <0.05. **d**, Ranking of cell-type relevance to diseases based on heritability enrichment. *Regression coefficient*, from the analysis in **(c)**, can be interpreted as the extent of cell-type specific heritability enrichment, and thus cell-type relevance. *Error bars*, standard error of the estimate. *Blue line* and *grey shade*, linear regression mean and 95% confidence intervals. **e**, Mapping disease-associated variants to candidate genes using CREs with cell-type and cell-state contexts. *Top*, percentage of loci with at least 1 candidate gene mapped. *Middle* and *bottom*, number of candidate genes mapped using proximal and distal CREs, respectively, with cell-type or cell-state contexts. **f**, Genetic signals and tCREs at a Crohn’s disease risk locus close to *PTGER4*. Crohn’s disease associated SNP, in LD with r^2^ ≥0.5, represented by the height of the bars. Co-activity among tCREs, with co-activity score ≥0.2, represented by the color of the arcs. Resting and stimulated PBMC data were pooled in the sc-ATAC-seq and sc-end5-dT tracks. The view was generated in the Zenbu genome browser with modifications (Data availability). **g**, Cell-types and cell-states specific activity of proximal and distal tCREs of *PTGER4*. “*Merged distal tCREs*” refers to the sum of expression values of 20 closely located distal tCREs, as detailed in Supplementary Fig.13.

As we observed a generally higher level of heritability enrichment in distal tCREs than in distal aCREs (Fig. 6a, *bottom row*), we reasoned that transcription at CRE could be indicative of its activity and biological relevance. We therefore investigated the heritability enrichment in aCREs with various levels of transcription evidence (Fig. 6b). About 45% of all aCREs showed evidence of transcription (transcribed aCRE, Fig. 6b, *top row, right panel*). This percentage is comparable to our estimate that ∼47% of aCREs are transcribed in DMFB based on a bulk-CAGE dataset^47^ at an unprecedented sequencing depth of 12,000M reads (Supplementary Fig. 11; Methods), suggesting this percentage of transcribed aCRE in PBMC is a reasonable estimate despite limited sequencing depth at ∼1,000M reads (Supplementary Fig. 1a). Untranscribed aCREs may be poised promoters, untranscribed enhancers, silencers, insulators or technical artifacts of sc-ATAC-seq^18–20^. These untranscribed aCREs are not enriched in heritability for most traits, in contrast to the transcribed aCREs which showed significant heritability enrichment (Fig. 6b, *top row, left panel*, FDR <0.05). The enrichment levels of transcribed aCREs are dependent on the level of transcription, particularly in distal aCREs, where only ∼15% of which showed high evidence of transcription and are highly enriched in heritability (Fig. 6b, *bottom row*). These observations are consistent with previous reports^2,24^ and highlight the importance of considering the evidence of transcription to identify active enhancers.

We next examined the enrichment of heritability in cell-type-specific CREs, which may be used to identify trait-relevant cell-types (Fig. 6c,d; Methods). As expected, immune cell-type-specific CREs are not enriched in heritability of psychiatric and metabolic traits. Also, monocyte count heritability is enriched in monocyte-specific CREs and leukocyte count heritability is enriched in CREs specific to most cell-types (Fig. 6c, *hematologic panel, solid dots*, FDR <0.05). Investigating the heritability of immunologic disorders, we found consistent and significant enrichment of T-cell, B-cell and NK cell-specific CREs in most disorders (FDR <0.05), recapitulating the general relevance of lymphoid cells in these disorders^48^. While the sensitivities of tCRE and aCRE in detection of heritability enrichments are generally comparable in most disorders (Fig.6c, *solid dots*), we observed a slightly higher sensitivity in aCRE in some disorders, such as SLE and rheumatoid arthritis. Next we compared the extent of cell-type-specific enrichment of heritability^45^ in tCRE and aCRE as a metric to prioritize cell-type relevance for each disorders (Fig. 6d). We found an overall consistent cell-type ranking between tCRE and aCRE (mean Pearson’s r=0.61). Particularly, in Eczema with Pearson’s r of 0.90, both tCRE and aCRE consistently ranked CD4+ T-cells as the most relevant cell-type, recapitulating the pivotal roles of Type-1 and -2 immune responses in skin inflammation^49^. We have also performed the same analyses for stimulation-responsive CREs in various cell-types, with similar conclusions (Supplementary Fig. 12). In summary, our data demonstrates the applications of tCRE in the prioritization of trait-relevant cell-types.

### Functional annotation disease-associated variants using tCRE

Lastly, we compared the application of tCRE and aCRE in functional annotation of disease-associated variants by linking to their target genes in relevant cell-types (Methods). Using tCREs, ∼41% of the disease -associated loci could be connected to a relevant cell-type-specific CRE, compared to ∼68% by aCRE (Fig. 6e). In addition, we found the number of genes associated by distal CRE is ∼4.5 times lower in tCRE than aCRE. As the number of distal aCRE (n=129,679) is much larger than distal tCRE (n=26,266), the higher number of genes associated by distal aCREs is not surprising. However, given the lack of heritability enrichment in distal aCRE with no (∼62%) or low (∼23%) transcription evidence (Fig. 6b), and the generally lower validation rate of aCRE co-activity links (Fig. 5c), the relevance of the genes associated by these untranscribed distal aCREs remains elusive, despite the high number. Here we highlight Prostaglandin E2 receptor 4 (*PTERG4*) gene located in proximity to the linkage disequilibrium (LD) block associated with multiple sclerosis, allergy, asthma, Crohn’s disease and ulcerative colitis (Fig. 6f). We found a cluster of distal tCREs within these LD blocks (Supplementary Fig. 13), overlapping with multiple disease-associated variants and are linked by co-activity to the proximal tCREs of *PTERG4* (Fig. 6f). Both distal and proximal tCREs of *PTERG4* are highly enriched in T-cells, agreeing with the pivotal roles of T-cells in autoimmune disorders^50^ (Fig. 6g). These findings are consistent with a previous report demonstrating that the distal CREs found in Crohn’s disease risk locus might regulate the expression of *PTGER4*^51^. In summary, these observations demonstrated the applications of single-cell tCRE activities in functional annotation of disease-associated variants with epigenomic and cellular contexts. These data can be browsed online under “Variants Interpretations” (Data availability).

## Discussion

Here we presented an analytical framework using sc-end5-seq data to define transcribed CRE in single cells, for interrogating gene regulation and disease heritability with cell-type-specific context. Compared to accessibility data which is close-to-binary in nature^17^, transcription data is quantitative^23^ and has a wider dynamic range. This explains the higher accuracy in predicting CRE interactions by co-activity by tCRE (Fig. 5e). In addition, the dynamic nature of transcription might better capture the granularities of gene regulation during rapid cell-state changes, as reflected in the detection of shifts in alternative promoter usage by transcription data, but not by accessibility data (Fig. 5h). The lack of heritability enrichment in untranscribed aCREs (Fig. 6b) and higher levels of heritability enrichment in distal tCRE (Fig. 6a), also highlight the importance of considering the evidence of transcription to identify active and biologically relevant CREs. Although we demonstrated that sc-end5-seq methods can detect enhancer RNAs (Fig. 4h), the high level of dropouts (due to their low abundance) renders the analyses of enhancer RNAs in single cells challenging. One might partially alleviate the problem by pooling data from multiple similar cells (as metacells) for downstream analyses. Alternatively, constructing the sc-end5-seq libraries with nuclei instead of whole cells^52^ or targeted capturing of a subset of enhancer RNAs^53^, should enrich enhancer RNAs in the library to reduce dropouts. Currently, most datasets generated on the Chromium platform (10x Genomics) are from sc-end3-dT, while the sc-end5-dT method is used only when T-or B-cell receptor repertoires are considered. Although it is well-known that sc-end5-seq data can theoretically detect CRE activity at no extra cost, the lack of dedicated tools for data analyses, in particular *de novo* CRE discovery, prevented the wider adoption of this analytical framework. Here we presented *SCAFE* for dedicated analyses of transcribed CREs (Supplementary Fig. 4) and we anticipate wide applications of sc-end5-seq methods along with *SCAFE* in the future for interrogating CREs in single cells.

## Methods

### Human ethics

All human samples examined in this study were either exempted material or were obtained with informed consent and covered under the research protocol (no. H30-9) approved by the ethics committees of the RIKEN Yokohama Campus.

### Genome version and gene models

Human genome assembly version hg19 and gene models from GENCODE^55^ version v32lift37 were used in all analyses of this study, unless otherwise stated.

### Preparing DMFB and iPSC samples

DMFB from neonatal foreskin were purchased (Lonza). Cells were cultured in Gibco Dulbecco’s Modified Eagle Medium (DMEM, high glucose with L-glutamine) supplemented with 10% Fetal bovine serum (FBS) and penicillin/streptomycin. Cells were dissociated with trypsin in 0.25% Ethylenediaminetetraacetic acid (EDTA) for 5 minutes at 37°C and then washed twice with 0.04% Bovine serum albumin (BSA) in Phosphate-buffered saline (PBS). iPSC^56^ were cultured in StemFit medium (Reprocell) under feeder-free conditions at 37°C in a 5% CO_2_ incubator. The cells were plated on dishes pre-coated with iMatrix-511 (Nippi). Rock inhibitor Y-27632 (FUJIFILM Wako) was added to the cells at a final concentration of 10μM during the first day of culturing. StemFit medium is refreshed daily until harvesting. The cells were detached and dissociated by incubating with TrypLE Select (Thermo Fisher Scientific) followed by scrapping in StemFit medium. The cells were collected by centrifuge and washed twice with 0.04% BSA in PBS.

### Preparing PBMC samples

Human PBMCs were prepared from the whole blood freshly collected from a male healthy donor with Leucosep (Greiner). Isolated 2×10^6^ PBMC cells were incubated with PMA/ionomycin (i.e., stimulated) (Cell Activation Cocktail with Brefeldin A, Biolegend), or DMSO as control (i.e., resting), for six hours.

### Isolating cytoplasmic, nucleoplasmic, and chromatin-bound RNAs

Subcellular fractionation of DMFB and iPSC cells was carried out according to a previous study^57^. Briefly, cells were grown to ∼90% confluency in 10cm dishes were collected by trypsinization and washed once with 0.04% BSA in PBS. The cells were lysed in lysis buffer containing 10 mM Tris pH 7.4, 150 mM NaCl and 0.15% IGEPAL CA-640 (Sigma-Aldrich), followed by separation of the nuclei from the cytoplasmic materials by centrifugation in a sucrose cushion containing 10 mM Tris pH 7.4, 150 mM NaCl and 24% sucrose at 1,000 × g for 10 minutes at 4ºC. The isolated nuclei were rinsed once in PBS-EDTA and lysed by adding glycerol buffer (20 mM Tris pH 7.4, 75 mM NaCl, 0.5 mM EDTA and 50% glycerol) and urea buffer (10 mM Tris pH 7.4, 1 M urea, 0.3 M NaCl, 7.5 mM MgCl_2_, 0.2 mM EDTA and 1% IGEPAL) in equal volumes. The precipitate, which contained the chromatin-RNA complex, was isolated by centrifugation at 13,000 × g for 2 minutes at 4ºC and washed once in PBS-EDTA. RNA from each of the three subcellular fractions was isolated by TRIzol Reagent (Thermo Fisher Scientific) according to the manufacturer’s instructions.

### Preparing bulk-CAGE, bulk-RNA-seq and bulk-ATAC-seq libraries for DMFB and iPSC

Bulk-CAGE libraries were generated by the nAnT-iCAGE^58^ method as previously described and sequenced on HiSeq 2500 (Illumina) as 50bp single-end reads. Bulk-RNA-seq libraries was generated as previously described^2^ and sequenced on HiSeq 2500 (Illumina) as 100bp paired-end reads. Bulk-ATAC-seq was performed as previously described^59^ with slight modifications. Briefly, 25,000 cells were used for library preparation. Due to the more resistant membrane properties of DMFB cells, 0.25% IGEPAL CA-630 (Sigma-Aldrich) were used for cell lysis. Transposase reaction was carried out as described followed by 10 to 12 cycles of PCR amplification. Amplified DNA fragments were purified with MinElute PCR Purification Kit (QIAGEN) and size-selected with AMPure XP (Beckman Coulter). All libraries were examined in Bioanalyzer (Agilent) and quantified by KAPA Library Quantification Kits (Kapa Biosystems). Bulk-ATAC-seq libraries were sequenced on HiSeq 2500 (Illumina) as 50bp paired-end reads.

### Preparing sc-RNA-seq libraries for DMFB and iPSC

Freshly prepared iPSC and DMFB cells were loaded onto the Chromium Controller (10x Genomics) on different days. Cell number and viability were measured by Countess II Automated Cell Counter (Thermo Fisher Scientific). Final cell density before loading was adjusted to 1.0×10^6^ cells/ml with >95% viability, targeting ∼5,000 cells. For sc-end3-dT libraries, we used Chromium Single Cell 3′ Reagent Kit v2 (10x Genomics). Briefly, single cells suspensions were mixed with the Single Cell Master Mix containing template switch oligonucleotide (AAGCAGTGGTATCAACGCAGAGTACATrGrGrG) and loaded together with 3′gel beads and partitioning oil into a Chip A Single Cell according to the manufacturer’s instructions (10x Genomics). For sc-end5-dT and sc-end5-rand libraries, we used Chromium Single Cell 5’ Library and Gel Bead Kit (initial version, 10x Genomics). Briefly, single cell suspensions were mixed with Single Cell Master Mix containing oligo(dT) primer (AAGCAGTGGTATCAACGCAGAGTAC– T(30)–VN) or random primer (17 µM; AAGCAGTGGTATCAACGCAGAGTACNNNNNN) as RT primer, and loaded together with 5′gel beads and partitioning oil into a Chip A Single Cell according to the manufacturer’s instructions. RNAs within single cells were uniquely barcoded and reverse transcribed within droplets. Both methods used Veriti Thermal Cycler (Applied Biosystems) for RT reaction at 53ºC for 45 minutes. Then, cDNAs from each method were amplified using cDNA primer mix from the kit, with 12 and 14 PCR cycles for sc-end5-dT and sc-end5-rand respectively, followed by the standard steps according to manufacturer’s instructions. For iPSC and DMFB, six libraries (i.e., 3 methods × 2 cell lines) were barcoded by different indexes from i7 sample index plate (10x Genomics). The libraries were examined in Bioanalyzer (Agilent) for size profiles and quantified by KAPA Library Quantification Kits (Kapa Biosystems). All libraries were sequenced on HiSeq 2500 (Illumina) as 75 bp paired-end reads.

### Preparing sc-end5-dT and sc-ATAC-seq libraries for PBMC

Freshly prepared resting and stimulated PBMCs were subjected to sc-end5-dT (Chromium Single Cell 5’ Library and Gel Bead Kit, initial version) and sc-ATAC-seq (Chromium Single Cell ATAC Reagent Kit v1) library construction on the same day using the Chromium platform according to manufacturer’s instructions (10x Genomics). About 5,000 cells/nuclei were targeted per reaction. sc-end5-dT and sc-ATAC-seq libraries were sequenced on HiSeq 2500 (Illumina) as 75bp and 100bp paired-end reads respectively.

### Processing bulk-RNA-seq and bulk-CAGE data for DMFB and iPSC

Reads were aligned to hg19 with *hisat2 v2*.*0*.*4*^60^ using default parameters. For each sample, the first aligned base at the 5’end of read 1 was piled up to a CTSS (i.e., Capped-TSS) bed file using custom *perl* scripts, available at https://github.com/chung-lab/scafe. These CTSS bed files were used for down-sampling, feature intersection and counting. Bulk-RNA-seq and bulk-CAGE data were processed with the same procedures.

### Processing FANTOM6 bulk-CAGE data for DMFB

Publicly available bulk-CAGE dataset on DMFB (n=1,163) were obtained^47^. Alignment bam files (on hg38) were converted to CTSS bed files as described above and lifted over to hg19 using *liftover* (http://genome.ucsc.edu). All CTSS bed files were pooled and subsampled to various depths. These subsampled CTSS bed files were processed in the *SCAFE* for *de novo* definition of TSS clusters and calculation of their logistic probabilities as described below.

### Processing bulk-ATAC-seq data for DMFB and iPSC

The bulk-ATAC-seq data for DMFB and iPSC were processed using ENCODE consortium pipelines (https://github.com/kundajelab/atac_dnase_pipelines). The – log(P) signal tracks for pooled replicates were used to defined gold-standards for training of the TSS classifiers.

### Processing sc-RNA-seq data for DMFB and iPSC

Reads were aligned to hg19 with *Cell Ranger v3*.*1*.*0* (10x Genomics), and bam files were processed with *SCAFE* to generate filtered CTSS bed files and *de novo* define tCRE. Annotation counts were produced by intersecting CTSS bed files with GENCODE gene models. Metagene plots from overlapping CTSS bed files with exons binned with Bioconductor *equisplit* using *foverlaps*. Enrichment of genesets in sc-end5-dT versus sc-end5-rand was tested using *fgsea v1*.*16*.*0*^61^ with nperm=1000. Genesets were defined as: 1) cytoplasmic, nucleoplasmic, and chromatin-bound RNAs: log_2_ fold-change ≥2 in fractionated CAGE data compared to total CAGE data, 2) long and short RNAs: maximum transcript length per gene ≥25,000nt and <1,000nt, 3) Non-polyA histone RNAs: histone RNAs with log_2_ fold-change ≥2 in non-polyA fraction in a previous study^30^.

### Processing sc-end5-dT data for PBMC

Reads were aligned to hg19 with *Cell Ranger* and the gene-based expression matrixes were processed with *Seurat v3*^62^. Briefly, cells were excluded with ≥4 median absolute deviation from the mean for number of features, UMI count, and percentage of mitochondrial UMI. Top 2,000 variable features were selected. Resting and stimulated PBMC samples were integrated with Canonical correlation analysis (CCA) implemented in *Seurat* using principal component (PC) 1 to 20 based on gene-based expression matrix. Bam files were processed with *SCAFE* to generate filtered CTSS bed files and *de novo* define tCRE. tCRE-based expression matrices from *SCAFE* were added to the *Seurat* object for downstream analysis. Cell annotation was performed by manually combining annotations from *scMatch (version at 2020-10-10)*^63^ and known marker genes. Cell-type-specificity and stimulation-specificity of tCREs were calculated with *Seurat FindMarkers* function with min.pct=0, return.thresh=Inf, logfc.threshold=0, min.cells.group=0.

### Processing sc-ATAC-seq data for PBMC

Reads were aligned to hg19 with *Cell Ranger ATAC v1*.*2* (10x Genomics) and the data were processed with *SnapATAC v1*.*0*.*0*^64^ using default parameters, selecting cells with ≥40% reads in ATAC peaks. Resting and stimulated cells were integrated with *Harmony v1*.*0*^65^ using PC 1 to 20. sc-ATAC-seq and sc-end5-dT were integrated using *SnapATAC FindTransferAnchors* and *TransferData* functions to transfer cell cluster annotations from the sc-end5-dT cells to the sc-ATAC-seq cells. sc-ATAC-seq peaks were defined per cell-type using *SnapATAC runMACS*, then merged. These merged peaks were referred to as aCREs and these aCREs were annotated using *SCAFE*. Cell-type-specificity and stimulation-specificity of aCREs were calculated with *SnapATAC findDAR*.

### Estimating TF motif activity

*ChromVAR v1*.*12*.*0*^42^ was used to calculate per-cell TF motif activities for the JASPAR2018^66^ core motif set for tCRE or aCRE excluding chrM. The tCRE expression matrix was binarized prior to running. Fisher’s exact tests and correlations of the top 80 motifs by *ChromVAR* deviation score per cell-type were used in Supplementary Fig 9.

### Predicting CRE interaction by co-activity

*Cicero v1*.*3*.*4*.*11*^16^ was used to calculate the co-activity score between CRE pairs using default parameters. Only tCREs and aCREs present in ≥3 cells were considered. Co-activity scores were estimated separately using cells within individual cell-types (cell-type sets) or all cells (pooled set). A pair of CREs with co-activity score ≥0.2 is defined as “linked”. PCHi-C connections^43^ (without cutoffs) from all cell-types were pooled and used for validation of co-activity linked CREs pairs. For comparisons of validation rates between tCREs and aCREs, only a subset of CREs that are overlapped between tCREs and aCREs and CRE pairs located ≥10kb apart was used.

### Detecting shifts in alternative promoter usage

For each cell-type (excluding dendritic cells due to low cell count), knn clustering of the *Seurat* SNN matrix (k=50) was used to generate metacells. The proportion of UMI in each genes arising from proximal tCREs was calculated for each metacell. Cell-type-specific tCRE switching events were identified using a *t*-test for differences in the proportion of UMI in gene contributed from each tCRE between metacells of selected cell-type and a background of all other cell-types. sc-ATAC-seq signal (UMI per millions) at a tCRE was defined as the maximum signal in cell-type bigwig files generated with *SnapATAC runMACS*.

### Identifying 5′end of cDNA and unencoded-G

Previous studies suggest most reads derived from capped RNAs begin with an unencoded “G”, which can be used to distinguish genuine TSS from artifacts^26,67^. To precisely calculate the number of unencoded-G for each mapped read, we first identify the junction between TS oligo and cDNA sequence and then examine the cDNA 5′end. Specifically, to precisely locate the TS oligo-cDNA junction, we considered only the reads 1) containing the last 5nt of TS oligo sequence (i.e., ATGGG) with maximum one mismatch, 2) starting with a soft-clip region (“S” in CIGAR string^68^) of ±50% of the TS oligo sequence length (6nt to 20nt), 3) with a match region ≥5nt (“M” in CIGAR string) following the soft-clip region. The 5′end of cDNA was defined as the first nucleotide immediately following the last nucleotide of the TS oligo sequence. The first 3nt of cDNA sequence was compared to the genomic sequence at their corresponding aligned position, and the number of Gs that are mismatched was defined as the number of unencoded-G for the examined read.

### Removing strand invasion artifacts

Strand invasion artifacts, i.e., strand invaders, can be identified based on complementarity of genomic sequence upstream of the mapped reads to TS oligo sequence, according to a study^34^, see also 10x Genomics technical note (https://support.10xgenomics.com/permalink/3ItKYUsoESnDpnFNnfgvNT). Briefly, we extracted a 13nt genomic sequence immediately upstream of the 5′end of cDNA, then globally aligned to the TS oligo sequence (TTTCTTATATGGG) and calculated the edit distance. A read is considered as an artifact of strand invasion when 1) the edit distance ≤5 and two of the three nucleotides immediately upstream were guanosines (Supplementary Fig.5), based on the previously proposed thresholds^34^.

### Defining TSS clusters and extracting their properties

The 5′end of cDNA (i.e., TSS) were extracted as described above, deduplicated as UMI, piled-up and clustered with *Paraclu*^69^ using default parameters. Only the TSS clusters with total UMI ≥5 and summit UMI ≥3 were retained. The following properties were extracted for each TSS cluster: 1) cluster count, 2) summit count, 3) flank count, 4) corrected expression and 5) unencoded-G percentage. Cluster, summit and flank count refers to UMI counts within the cluster, at its summit, and within a region flanking its summit (±75nt). Corrected expression refers to an expression value relative to its local background, based on the assumption that the level of exon painting artefact is positively correlated with the transcript abundance. Specifically, if the summit of a TSS cluster is located within genic regions, it will be assigned to either exon or intron, in either sense or antisense strand of the corresponding gene, or otherwise assigned to intergenic, as its local background. All annotated TSS regions (±250nt) were masked from these local backgrounds. The density of UMI per nucleotide within each local background is calculated (i.e., local background density). The corrected expression of a TSS cluster is calculated as the ratio of the density of UMI within the region flanking its summit (±75nt) to the density of its local background. Unencoded-G percentage refers to the percentage of UMI within the cluster that has ≥1 unencoded-G.

### Building a TSS classifier

To combine multiple properties of TSS cluster into a single classifier, we used multiple logistic regression implemented in the *caret*^70^ R package. For training of this classifier, we defined a set of “gold standard” TSS clusters based on their ATAC-seq signal (as mean –log(P) within TSS cluster). Specifically, the top and bottom 5% of TSS clusters, ranked by their ATAC-seq signal, were defined as positive and negative gold standards, and used for training of the logistic models at 5-fold cross-validation. The resulting logistic probability was used as the TSS classifier. The performance of this TSS classifier, and its constituent metrics, is measured as AUC, using the top and bottom 10% of TSS clusters as positive and negative gold standards for testing. The cutoff of logistic probability at 0.5 is defined as the default threshold. All the TSS clusters in this study are filtered with this default cutoff (refer to as the “moderate” sets). In the PBMC datasets, corresponding sc-ATAC-seq datasets were used for training and an additional lenient logistic probability cutoff of 0.023 was used, which corresponds to a specificity of 0.5 (refer to as the “lenient” set).

### Defining tCRE and aCRE

tCREs are defined by merging closely located TSS clusters. Briefly, TSS clusters located within ±500nt of annotated gene TSS were classified as proximal, or as distal otherwise. All TSS clusters were then extended 400nt upstream and 100nt downstream. These extended ranges were merged using *bedtools*^71^, in a strand-specific manner for proximal TSS clusters and non-strand-specific manner for distal TSS clusters, as proximal and distal tCRE respectively. Distal tCRE were then assigned to either exonic, intronic or intergenic, in this order. aCREs are defined by the ATAC peak ranges output from *SnapATAC*. aCREs located within ±500nt of annotated gene TSS were classified as proximal, or as distal otherwise.

### Developing *SCAFE* tools

*SCAFE* (Single Cell Analysis of Five-prime Ends) consists of a set of command-line tools written in *perl* and *R* programming languages, for processing of sc-end5-seq data. Briefly, it takes a read alignment file (in bam format), maps the cDNA 5’ends, identifies genuine TSS clusters, defines tCREs, annotated tCREs to gene models, quantify their expression and predicts tCRE interactions by co-activity. The tools in *SCAFE* can be ran individually as independent tools or ran serially as predefined workflows. For details, please visit https://github.com/chung-lab/scafe.

### Processing of GWAS data

For heritability enrichment, GWAS summary statistics were obtained from (1) UK biobank heritability browser (https://nealelab.github.io/UKBB_ldsc/index.html) (for allergy, monocyte count, leukocyte count, autism, neuroticism, blood pressure) (2) Dr. Alkes Price group site (https://alkesgroup.broadinstitute.org/) (for Celiac disease, cholangitis, Crohn’s disease, eczema, rheumatoid arthritis (RA), systemic lupus erythematosus (SLE), ulcerative colitis, type 1 diabetes, blood glucose) and (3) Japanese encyclopedia of genetic associations (JENGER, http://jenger.riken.jp/) (for Grave’s disease). Summary statistics obtained from (1) and (2) were directly used for heritability enrichment analyses, while the summary statistics obtained from (3) were pre-processed using “*munge_sumstats*.*py”* scripts in *LDSC* software^72^. For linking trait-associated variants to candidate genes, lead variants (P <5×10^−8^) were obtained from (1) GWASdb^73^ (as of 19th August 2015, http://jjwanglab.org/gwasdb) and (2) NHGRI-EBI GWAS Catalog^74^ (release r2020-07-15). The variants within the LD block of these lead variants (i.e., proxy variants) were searched for using *PLINK v1*.*9*^75^ with an r^2^ ≥0.5 within ±500kb in matched population panels of Phase 3 1000 Genomes Project downloaded from MAGMA website^76^ (http://ctg.cncr.nl/software/MAGMA/ref_data/).

### Estimating enrichment of trait heritability

Enrichment of trait heritability in tCRE (or aCRE) was assessed by stratified LD score regression (S-LDSC) implemented in *LDSC* software. Briefly, sets of tCRE (or aCRE) were defined based on their proximity to annotated TSS (i.e., all, proximal or distal). Two sets of tCREs were used, including a default set and a lenient set as described in “Building a TSS classifier”. Additional sets of aCREs were generated based on evidence of transcription (i.e., number of UMI from sc-end5-dT reads as described in main text). Annotation files and LD score files were generated for each set of tCRE (or aCRE) using the “*make_annot*.*py*” and “*ldsc*.*py*” scripts using default parameters. Each set of tCRE (or aCRE) was added onto the 97 annotations of the baseline-LD model v2.2 and heritability enrichment (i.e., ratio of proportion of heritability to proportion of SNP) for each trait was estimated using the “*ldsc*.*py*” script with *“--h2”* flag in default parameters.

### Evaluating cell-type-specificity of trait heritability

Cell-type-specificity of trait heritability was assessed by LD score regression for specifically expressed genes (LDSC-SEG) implemented in *LDSC* software^77^. Briefly, enrichment of each tCRE (or aCRE) in each cell-type were calculated using *findDAR* implemented in *SnapATAC* and *FindMarkers* in *Seurat*, respectively. Sets of “cell-type-specific” tCRE (or aCRE) were defined as the top 20% of tCRE (or aCRE) ranked by the enrichment P for each cell-type. A set of “core” tCRE (or aCRE) was defined as all tCREs (or aCREs) that are not “cell-type-specific” to any of the cell-types. Annotation files and LD score files were generated for each set of “cell-type-specific” and “core” tCREs (or aCREs) using the “*make_annot*.*py*” and “*ldsc*.*py*” scripts using default parameters. For each cell-type, sets of “cell-type-specific” and “core” tCRE (or aCRE) were added onto the 53 annotations of baseline-LD model v1.2 and the contribution of “cell-type-specific” tCRE (or aCRE) to trait heritability (i.e., regression coefficient) for each trait was estimated using the “*ldsc*.*py*” script with *“--h2-cts”* flag in default parameters.

### Connecting trait-associated variants to candidate genes

Trait-associated variants were defined as mentioned above. A tCRE (or aCRE) is associated with a trait if it overlaps at least one trait-associated variant. A gene is associated with a trait when its proximal tCRE (or aCRE) is associated with a trait, or a distal tCRE (or aCRE) is associated with a trait and connected to its proximal tCRE by co-activity score ≥0.2.

### Interactive data analysis reports

Data can be interactively browsed as “analysis reports” in ZENBU^54^. These reports are organized into six themes: 1) TSS in Single Cells, 2) Benchmarking tssClusters, 3) Cell-type-specific CREs, 4) Alternative Promoter Usage, 5) Interactions between tCREs and 6) Variants Interpretations. In particular, the genome browser of ZENBU features on-the-fly demultiplexing single-cell or cell-type signals, enabling convenient and interactive interrogation of single-nucleotide resolution of transcription signal within each single cell. To explore the data, please visit https://fantom.gsc.riken.jp/zenbu/reports/#tCRE.report.index.

### Data visualization and statistics

We used the *ggplot2 R* package^78^ unless otherwise noted for visualizations.

## Data availability

Data are available in the ArrayExpress database (http://www.ebi.ac.uk/arrayexpress) under accession numbers: E-MTAB-10385 (sc-end5-dT, sc-end5-rand and sc-end3-dT for DMFB, iPSC), E-MTAB-10378 (sc-end5-dT for PBMC), E-MTAB-10381 (bulk-ATAC-seq for DMFB, iPSC), E-MTAB-10382 (sc-ATAC-seq for PBMC), E-MTAB-10383 (bulk-RNA-seq for DMFB, iPSC), E-MTAB-10384 (bulk-CAGE for DMFB, iPSC). These data can be interactively browsed on ZENBU^54^ at https://fantom.gsc.riken.jp/zenbu/reports/#tCRE.report.index.

## Code availability

The *SCAFE* software for processing 5′end RNA-seq data is available at https://github.com/chung-lab/scafe.

## Acknowledgements

This publication is part of the Human Cell Atlas -www.humancellatlas.org/publications. This research was supported by research grant to RIKEN Center for Integrative Medical Sciences from Ministry of Education, Culture, Sports, Science and Technology (MEXT). We would like to extend our thanks to GeNAS sequencing platform in IMS.

## Author contributions

CCH, JWS, PC conceived the project and supervised the research. TK optimized experiments and constructed single-cell libraries. JM and CCH analyzed most of the data. JM, TK, JWS, CCH wrote the manuscript. JCC processed the bulk-ATAC data. CWY processed the DMFB bulk CAGE data. CT and HS assisted the heritability enrichment analysis. AS, KY performed the PBMC stimulation experiments. YS performed cell fractionated bulk RNA experiments. JS assisted the building of ZENBU analysis reports. FLR performed bulk-ATAC-seq experiments. YA supported the logistics of sample collection.

---

**Supplementary Fig. 1.**
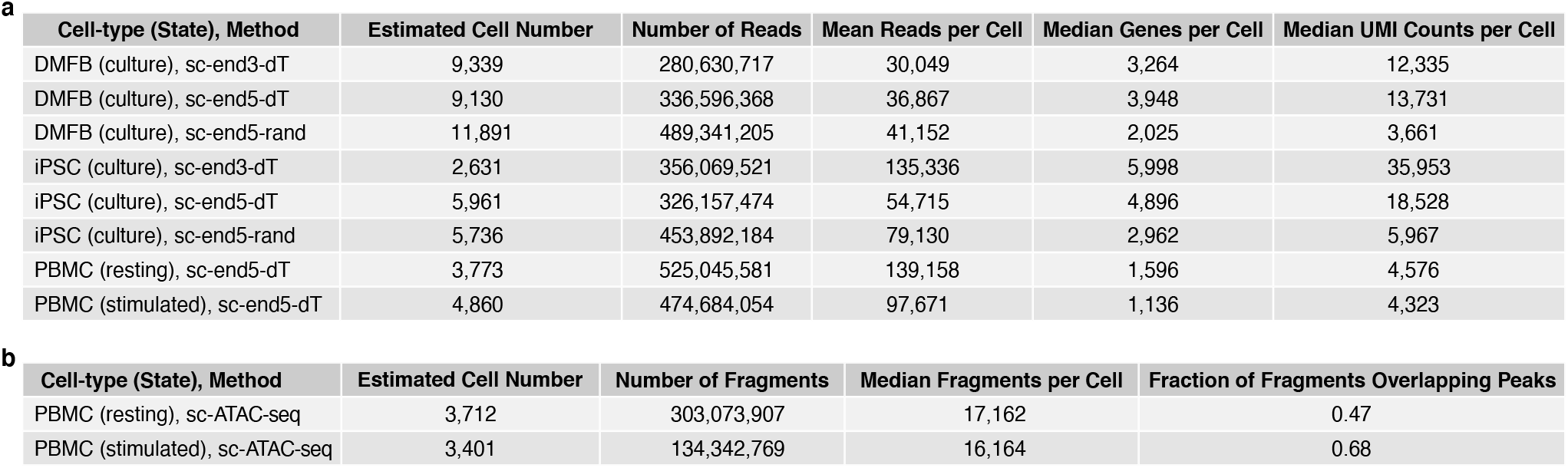
Statistics of sc-RNA-seq and sc-ATAC-seq libraries in this study. **a**, Statistics of sc-RNA-seq libraries. **b**, Statistics of sc-ATAC-seq libraries. All numbers were extracted from the reports generated from *Cell Ranger* (10x Genomics).

**Supplementary Fig. 2.**
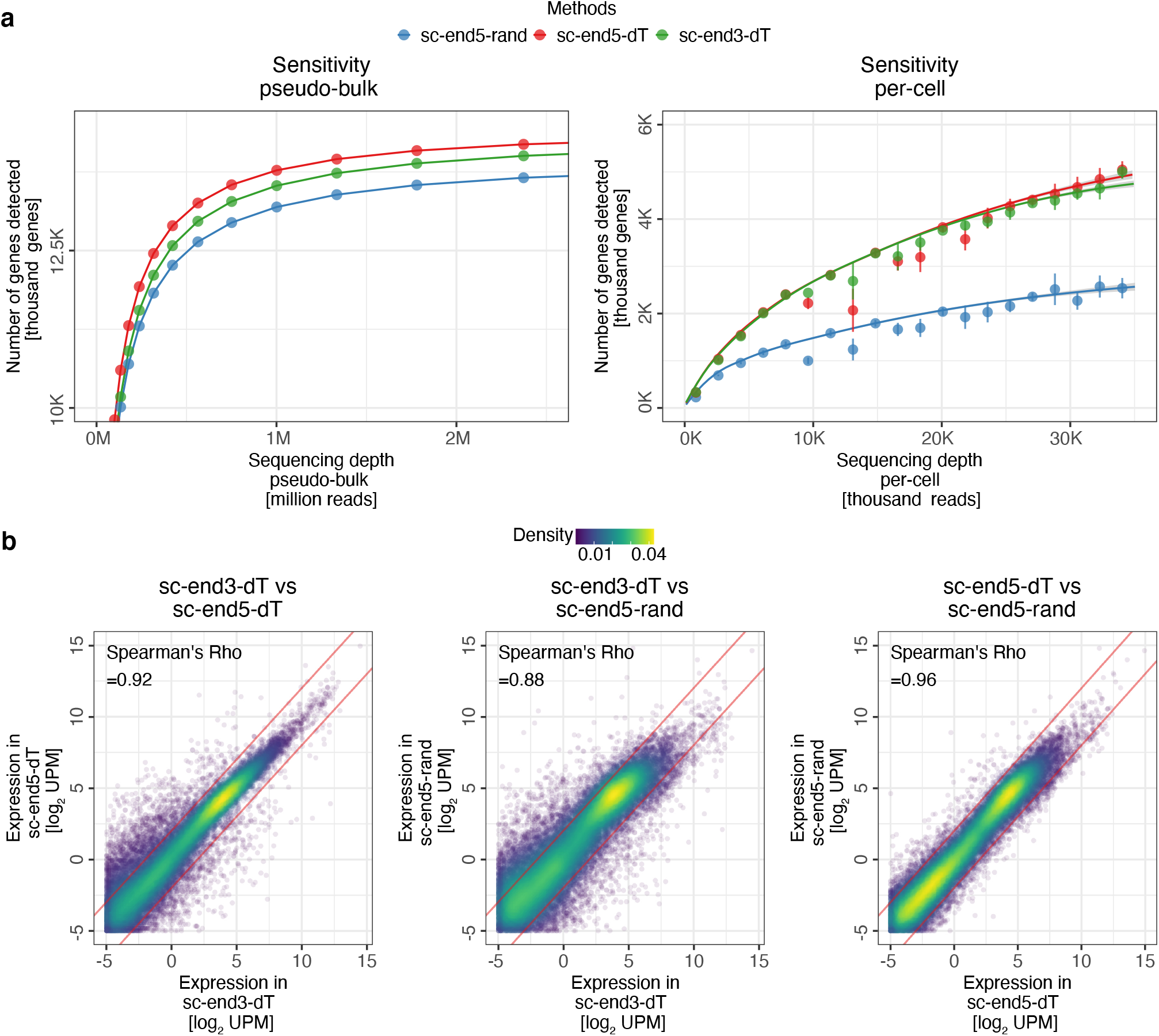
Performance of sc-RNA-seq methods in DMFB. **a**, Sensitivity of detection of genes in pseudo-bulk (*left*) and in single cells (*right*) across sequencing depth. *Error bars* represent standard deviation. The genes that are detected in bulk-RNA-seq were used to define the scope. **b**, Correlation of gene expression levels between the pseudo-bulk data of the three sc-RNA-seq methods. *Red line*, ±2-fold differences. *UPM*, UMI per million. Color represents the density of points.

**Supplementary Fig. 3.**
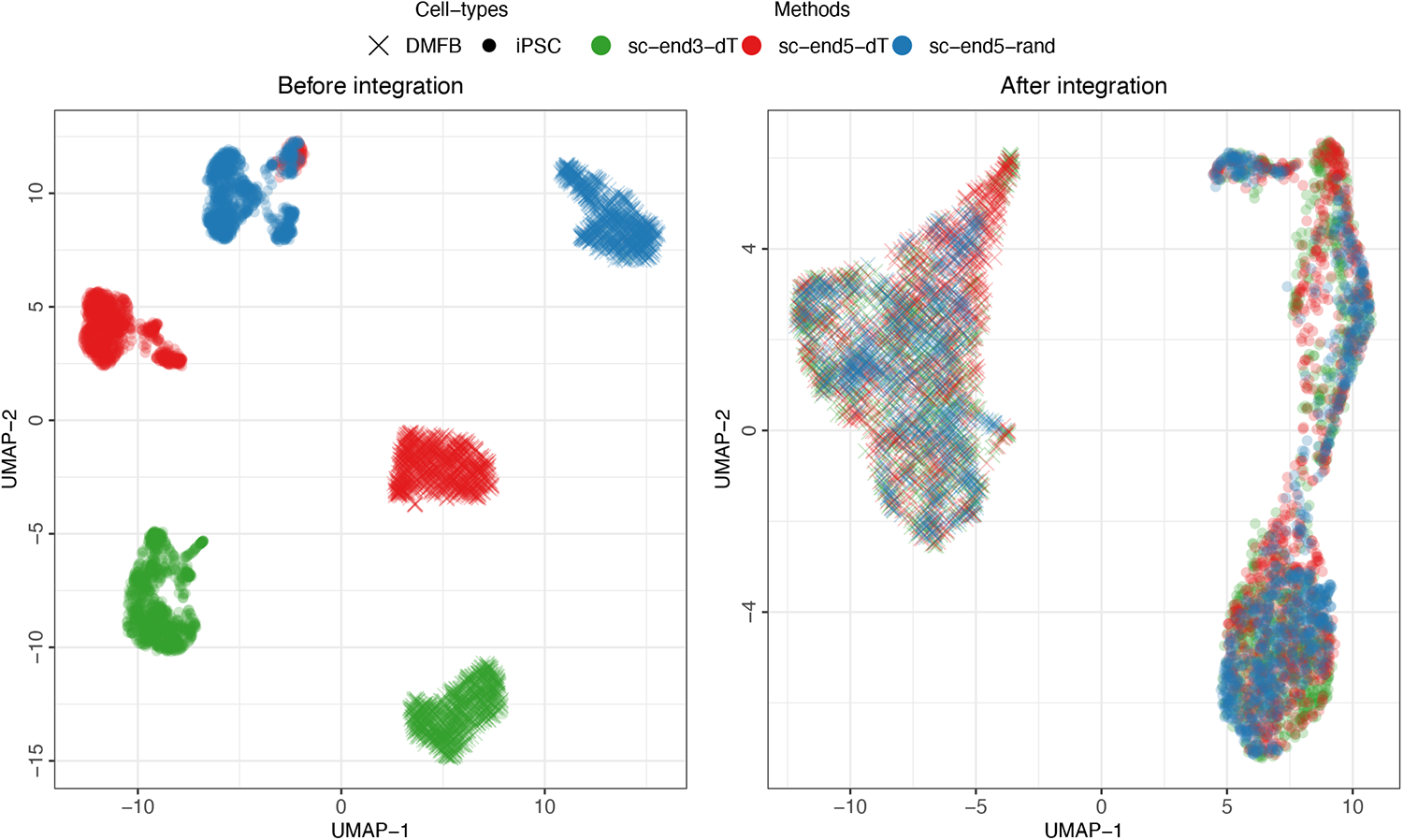
Integration of sc-RNA-seq methods. UMAP of sc-RNA-seq methods in iPSC and DMFB before integration (*left*), after *Seurat* CCA integration (*right*) demonstrating the ability to batch correct between different sequencing methods allowing the integration of sc-end5-seq datasets with existing sc-end3-seq resources.

**Supplementary Fig. 4.**
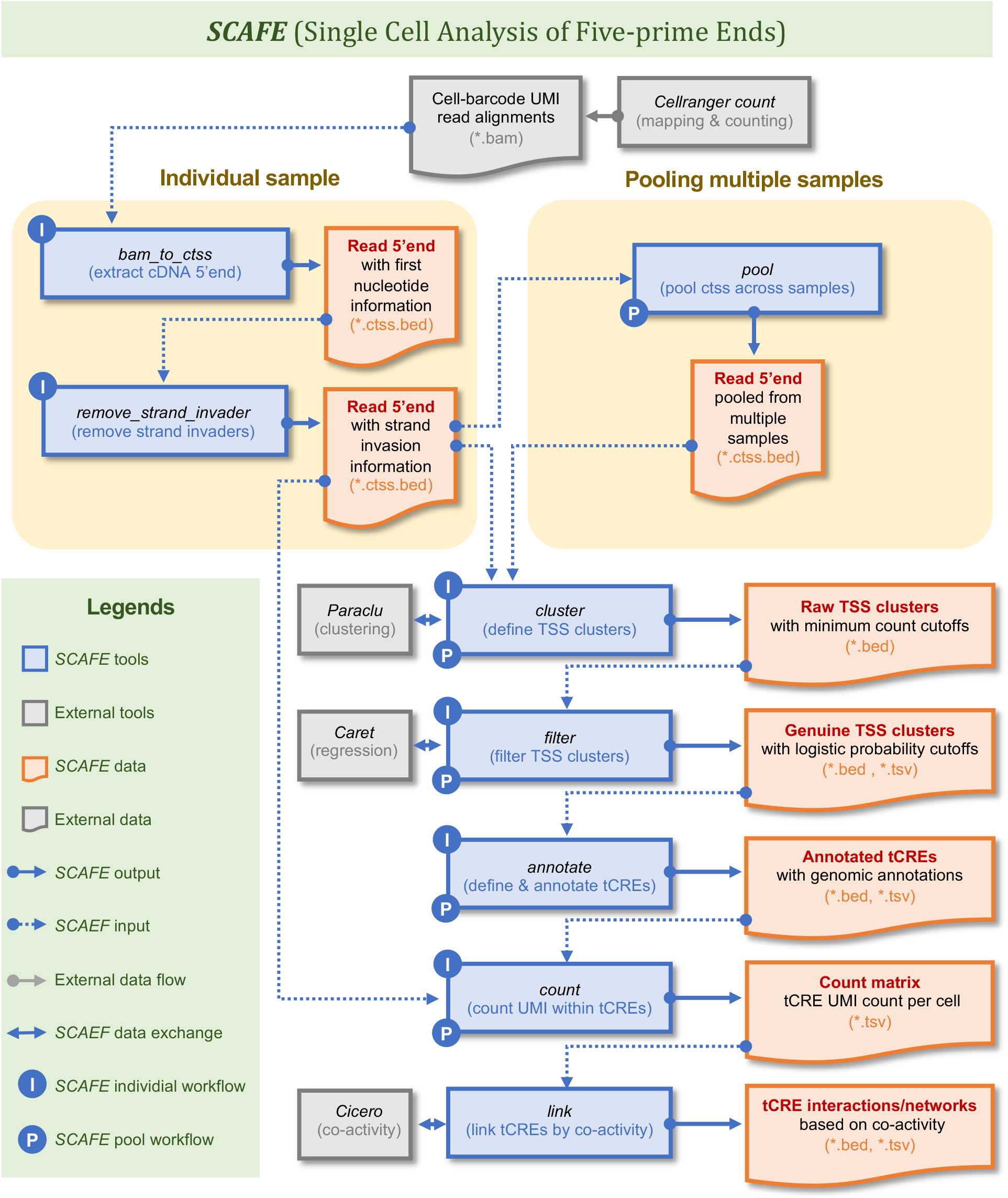
Overview of *SCAFE*. *SCAFE* consists of a set of *perl* programs for processing of sc-5end-seq data. Major tools are listed here, for all tools please visit https://github.com/chung-lab/scafe. *SCAFE* accepts read alignment in. *bam* format from *Cell Ranger* (10x Genomics). Tool *bam_to_ctss* extracts the 5′end of cDNA, taking the 5′ unencoded-Gs into account. Tool *remove_strand_invader* removes cDNA 5′ends that are strand invasion artifacts by aligning the TS oligo sequence to the immediate upstream sequence of the cDNA 5′end. Tool *cluster* performs clustering of cDNA 5′ends. Tool *filter* extracts the properties of TSS clusters and performs multiple logistic regression to distinguish genuine TSS clusters from artifacts. Tool *annotate* define tCREs by merging closely located TSS clusters and annotate tCREs based on their proximity to known genes. Tool *count* counts the number of UMI within each tCRE in single cells and generates a tCRE-Cell UMI count matrix. Tool *link* estimate the co-activity of tCRE pairs to predict CRE interaction and networks. *SCAFE* tools were also implemented workflows for processing of individual samples or pooling of multiple samples.

**Supplementary Fig. 5.**
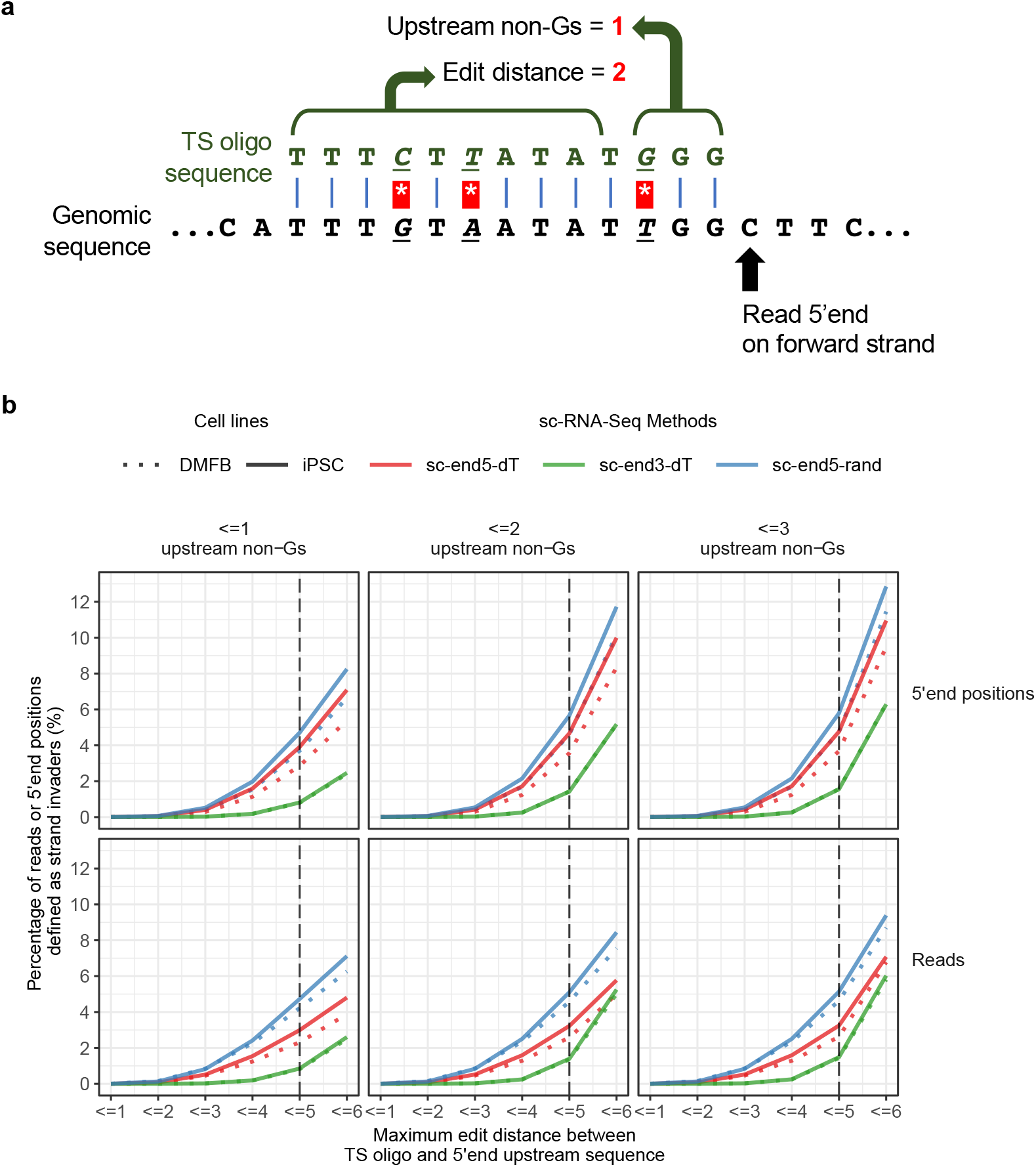
Detection of strand invasion artefacts. **a**, Rationale of strand invasion detection. The immediate upstream sequence of the read 5′end were aligned with TS oligo sequence. Number of upstream non-Gs was calculated from the first 3nt of the immediate upstream sequence. Edit distance was calculated from the last 10nt of the alignment. The shown example has 2 edit distances and 1 upstream non-Gs. **b**, Extent of strand invasion artefacts in various sc-RNA-seq methods. Maximum edit distance of 5 (*vertical dotted line*) and 2 upstream non-Gs (middle column) is chosen as the threshold to define strand invasion artefacts. At this threshold, the extent of strand invasion artefacts is consistently higher in sc-end5-rand (*blue*), compared to sc-end5-dT (*red*), in both DMFB and iPSC. sc-end3-dT (*green*) serves as a negative control of the random genomic background.

**Supplementary Fig. 6.**
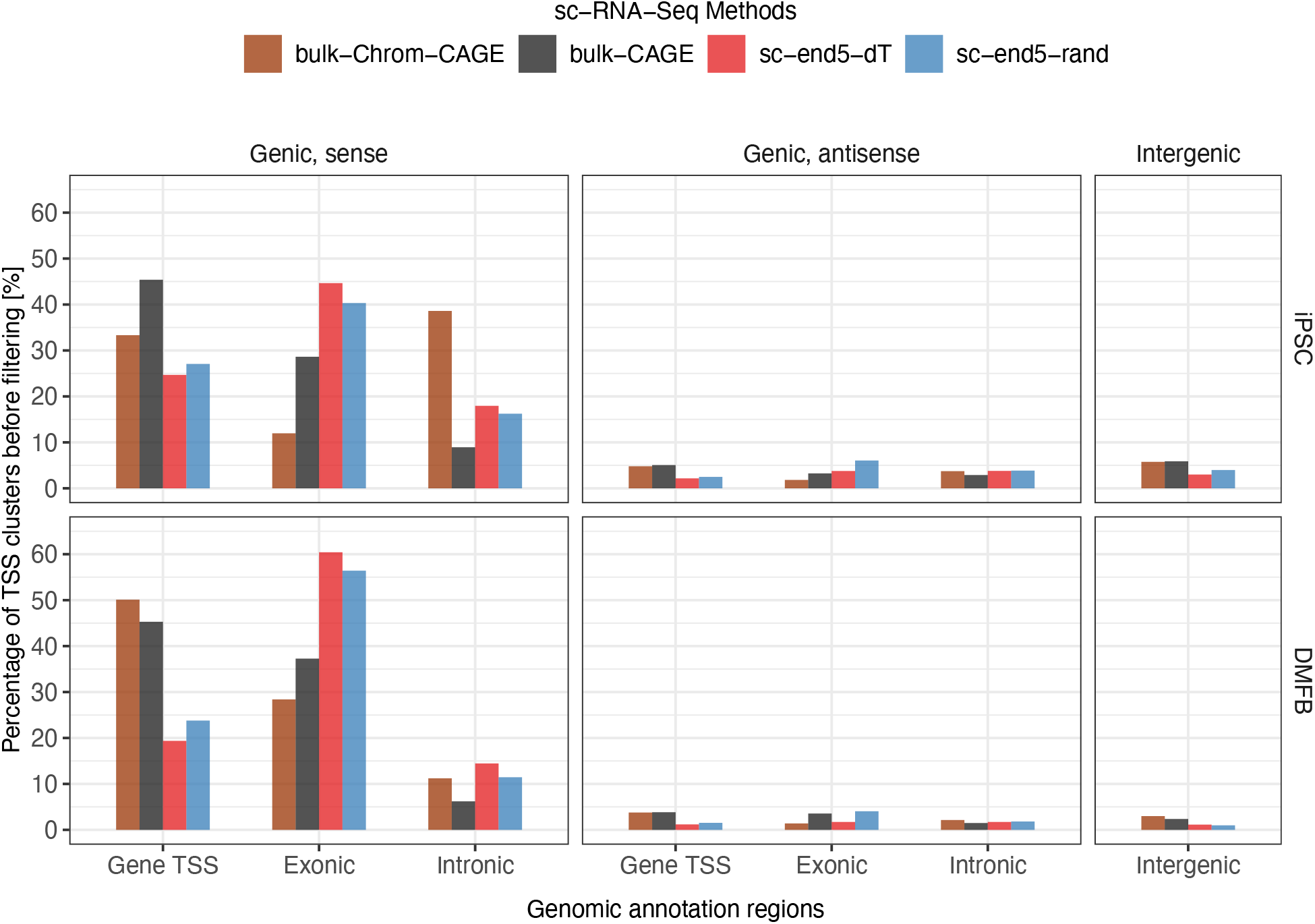
Genomic distribution of unfiltered TSS clusters. Unfiltered TSS clusters were assigned to various genic and intergenic annotations, based on their intersection with GENCODE annotation, in specific hierarchical orders (Methods). In both DMFB and iPSC, a large fraction of TSS clusters were assigned to the exonic and intronic regions in sense orientation, compared to that in the antisense orientation. This could be attributed to the “exon painting” artefacts as discussed, which could be filtered by considering various properties of the TSS clusters.

**Supplementary Fig. 7.**
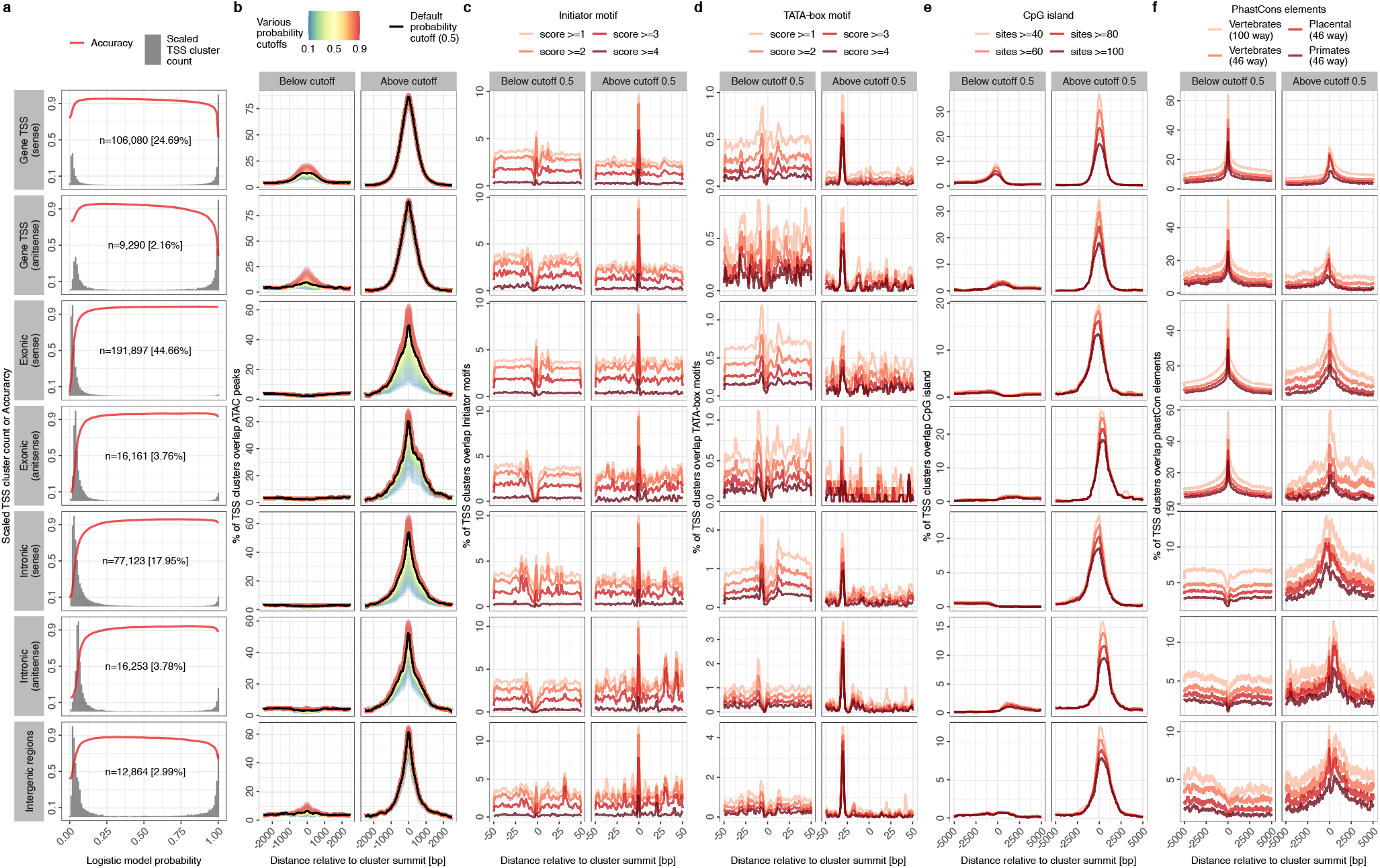
Properties of logistic model probability cutoffs for identification of genuine TSS clusters. **a**, Proportion of TSS clusters and accuracy along logistic model probability cutoffs. “n” and “%” refers to the number and percentage of TSS clusters in the category. **b**, Chromatin accessibility around summit of TSS clusters along logistic model probability thresholds. **c**,**d**,**e**,**f**, Distribution of Initiator motif, TATA-box motif, CpG island and PhastCons elements, respectively, around summit of TSS clusters below and above logistic model probability 0.5. Initiator motif and TATA-box motif were predicted on hg19 using *HOMER* (http://homer.ucsd.edu/homer/motif/). CpG island and PhastCons elements were downloaded from UCSC table browser (https://genome.ucsc.edu/). “Score” in *c* and *d* refers to score of motif prediction from *HOMER*. “Sites” in *e* refers to number of CG dinucleotides. In *f*, 100 ways and 46 ways refer to multiple alignments of 100 and 46 species respectively. Vertebrates, Placental and Primates refer to the scope of species used to define PhastCons elements. Initiator motif and TATA-box motif are, as expected, enriched at ∼0nt and ∼ –30nt, respectively, of the TSS cluster above below cutoff 0.5. The enrichment of PhastCons elements at the center of the “Gene TSS” and “Exonic” TSS clusters below cutoff 0.5 can be attributed to their overlap with exon regions, which are relative more conserved than intronic and intergenic regions.

**Supplementary Fig. 8.**
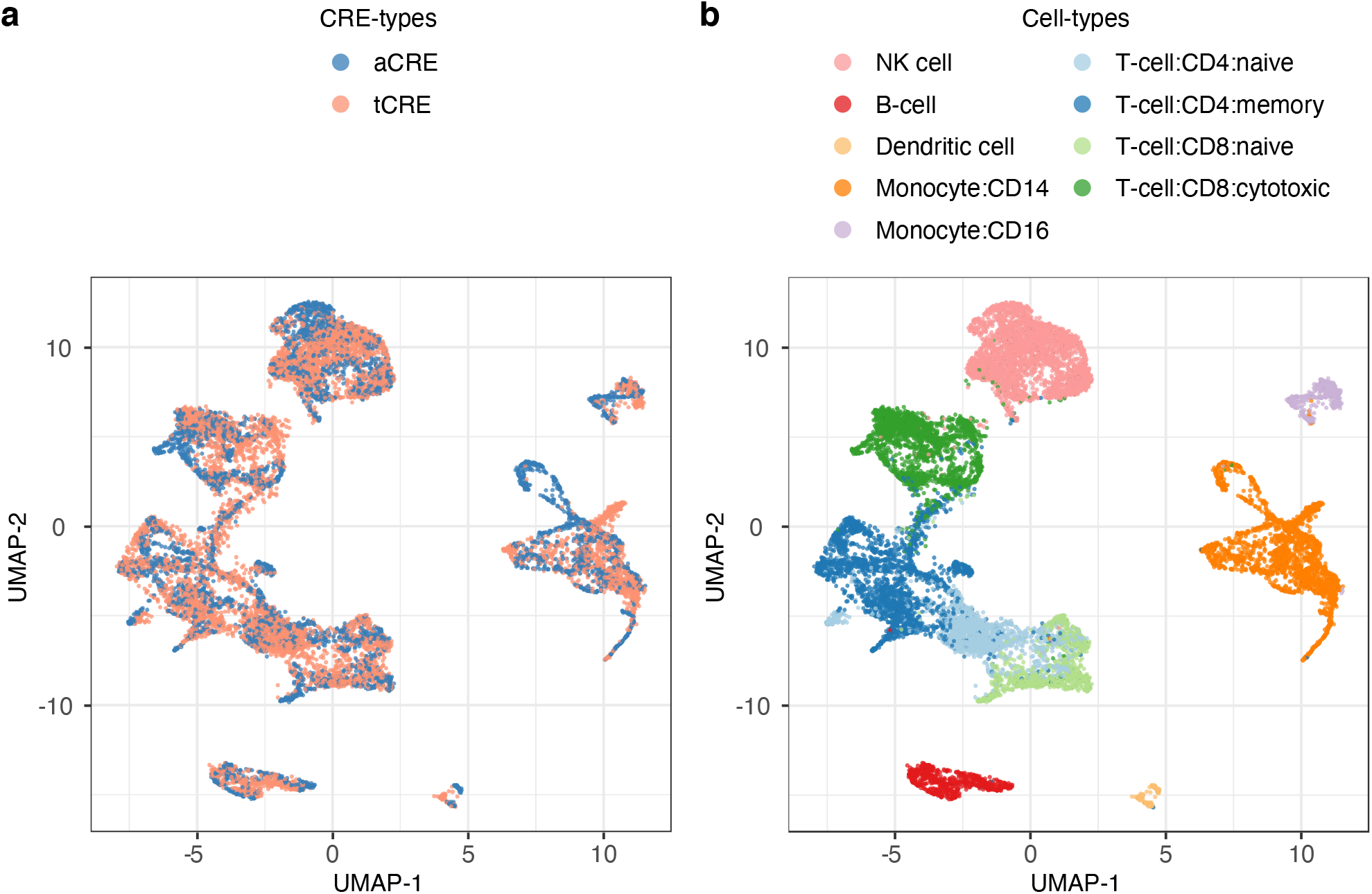
Integration of tCRE and aCRE. UMAP of tCRE and aCRE cells after integration by *Seurat CCA*. Colored by technology (*left*) and cell type annotation (*right*), cell type labels have been transferred from tCRE to aCRE.

**Supplementary Fig. 9.**
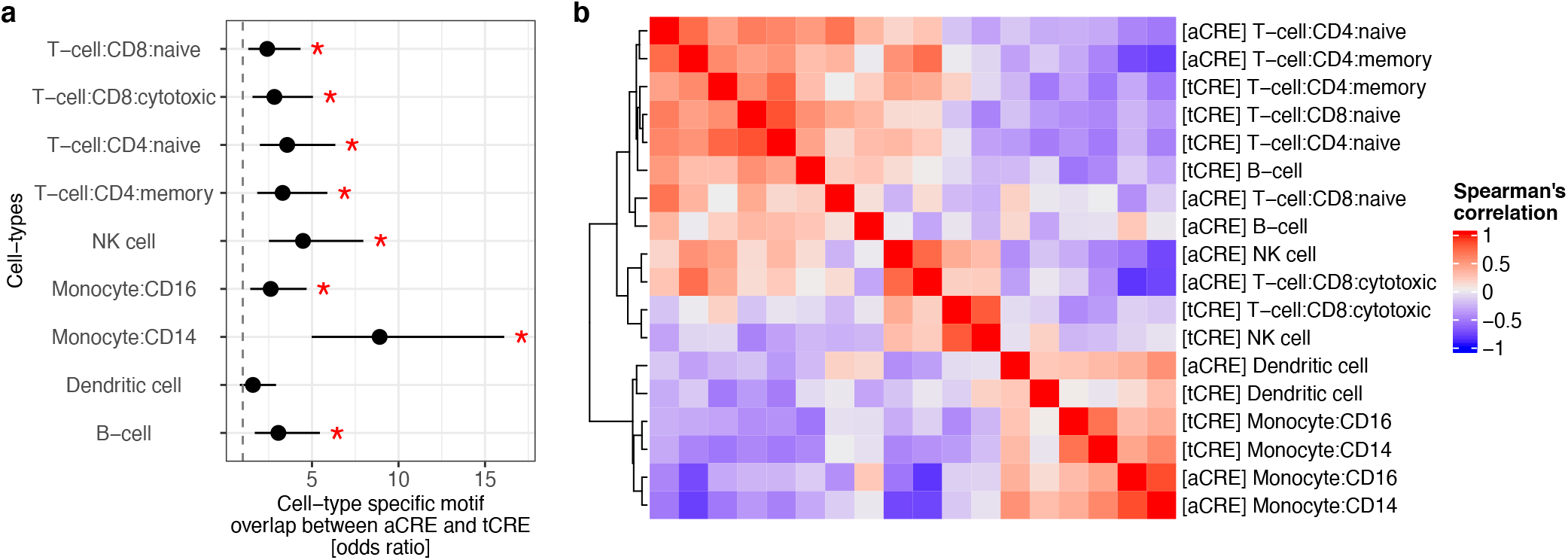
Cell-type specific TF motif activities based on tCRE and aCRE. **a**, Fisher’s exact test for odds ratio of overlap in top 80 cell-type specific TF motifs calculated with *ChromVAR* in aCRE and tCRE. *Asterisks*, P <0.05; *Error bars*, 95% confidence intervals; **b**, Heatmap of common cell-type specific motif activity from **(a)** averaged per cell-type (Spearman’s correlation).

**Supplementary Fig. 10.**
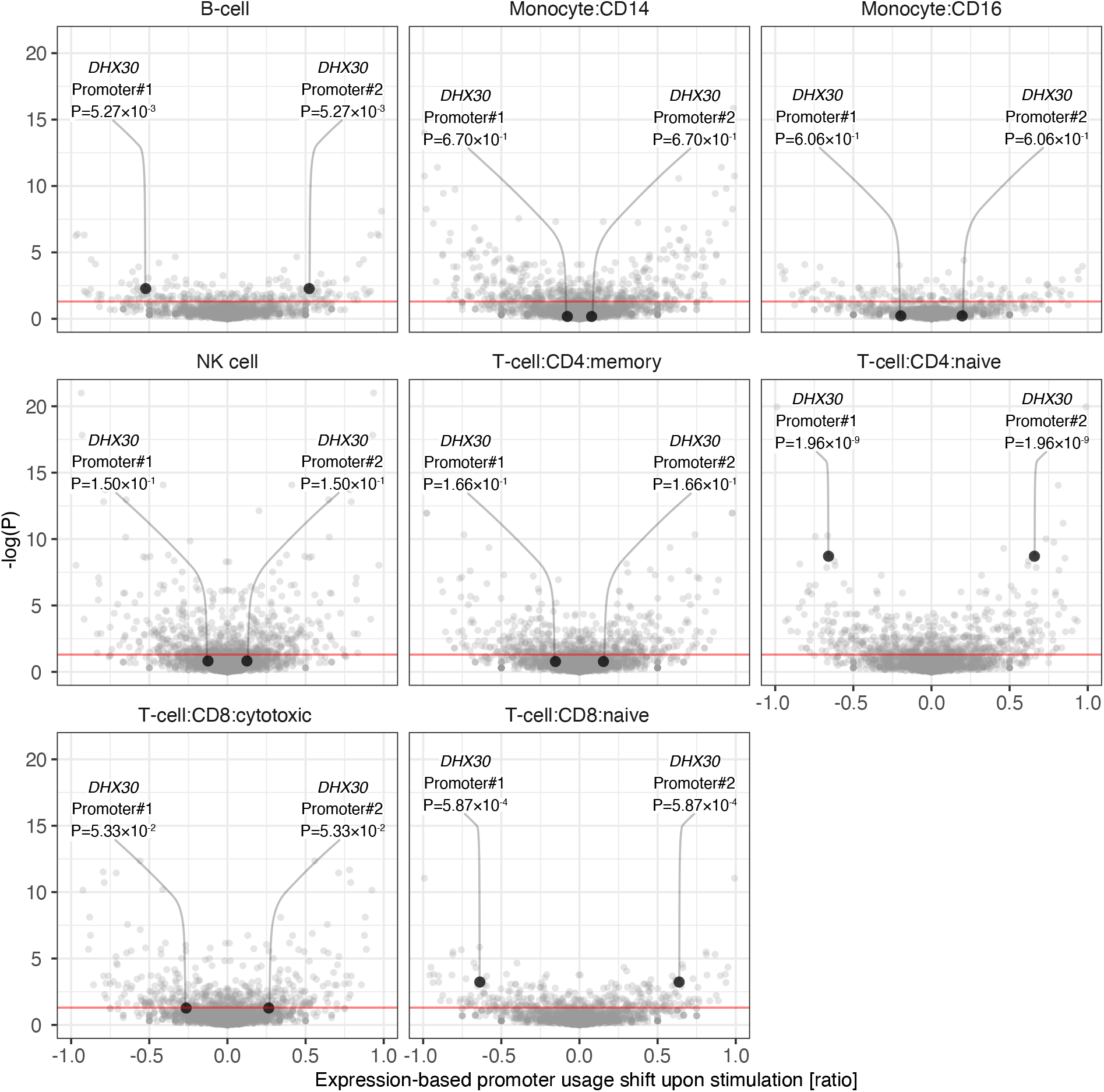
Cell-type specific shifts in alternative promoter usage. Each dot represents a proximal tCRE of a gene (i.e., promoter). Only genes with multiple alternative promoters are considered. *X-axis*, change in mean fraction of gene expression in metacells from each tCRE after stimulation. *Y-axis*, –log (P) of *t*-test for change in tCRE usage between metacells. Labeled example proximal tCREs of *DHX30* gene. *Red line*, P=0.05. Shift of usage from Promoter#1 to Promoter#2 occurs significantly (P <0.05) upon stimulation in T-cell:CD4:naïve, T-cell:CD8:naïve and B-cell.

**Supplementary Fig. 11.**
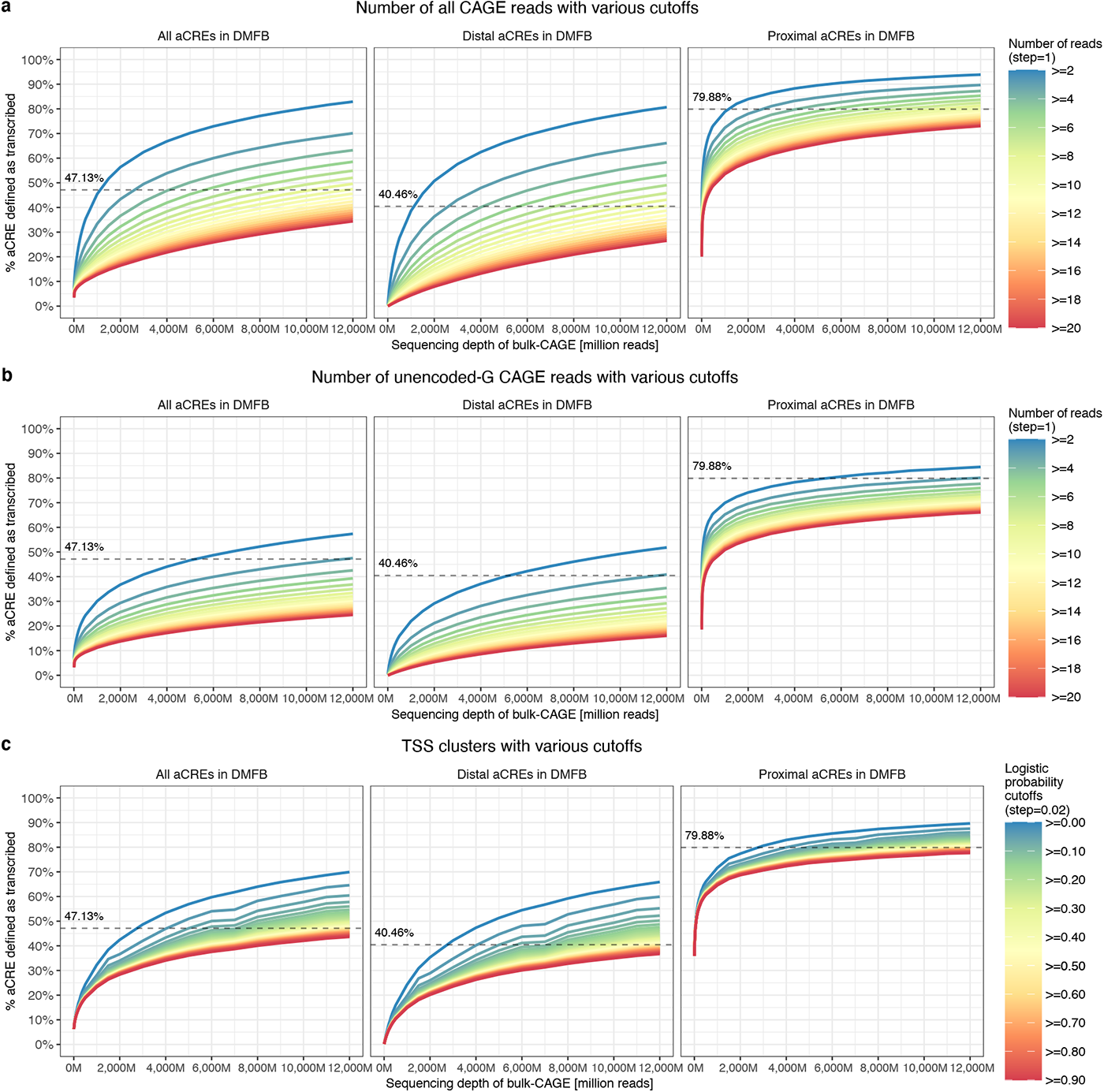
Percentage aCRE that are transcribed in DMFB. Estimating the percentage of aCREs that are transcribing using pooled CAGE libraries of DMFB at unprecedented sequencing depth based on **a**, number of all CAGE reads at TSS summit within aCRE, **b**, number of unencoded-G CAGE reads at TSS summit within aCRE, or **c**, highest logistic probability of TSS clusters within aCRE. *Dashed lines*, estimates of transcribed aCRE % at highest sequencing depth (i.e. 12,000M) based on TSS clusters with default logistic probability cutoffs at 0.5.

**Supplementary Fig. 12.**
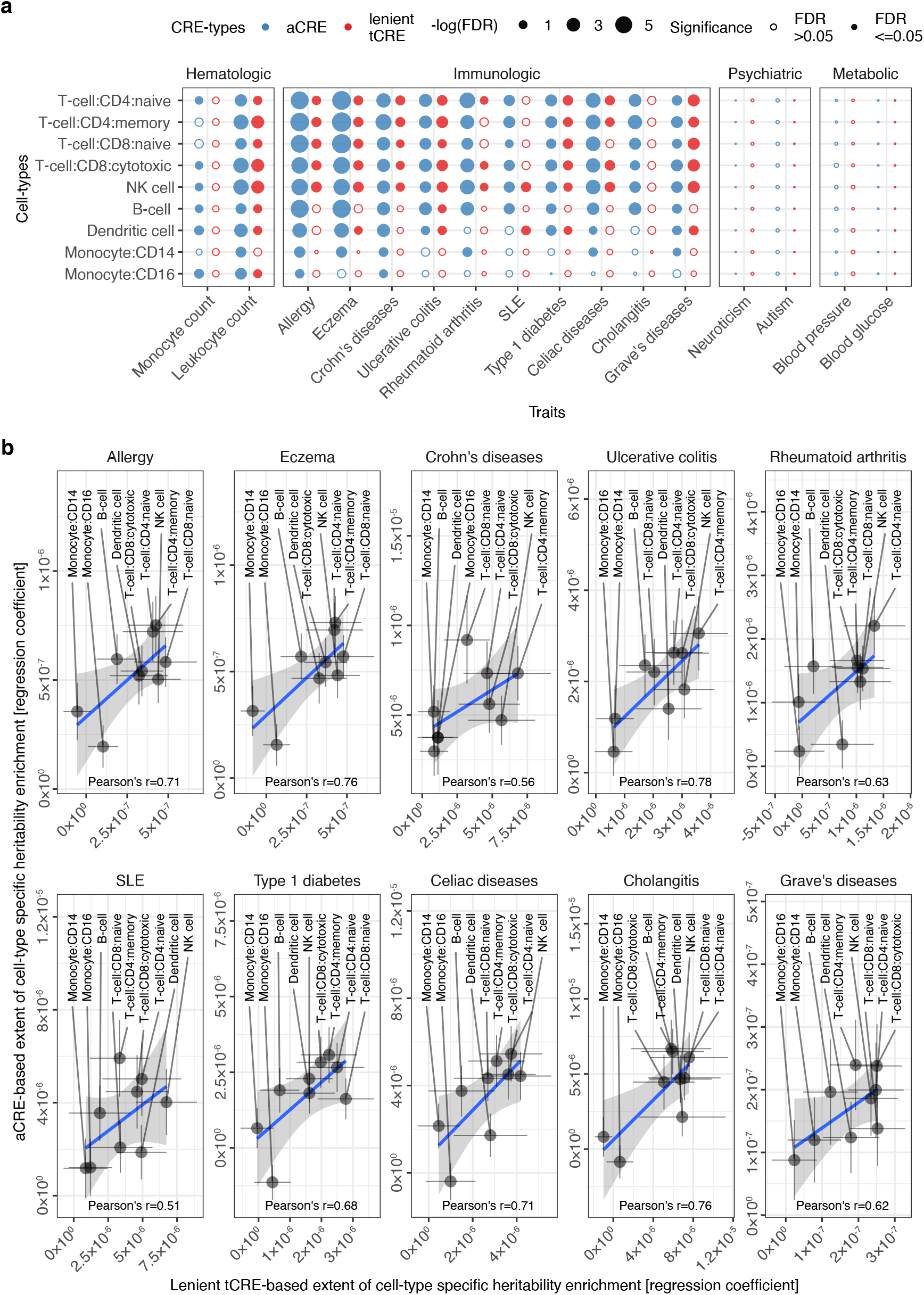
Heritability enrichment in stimulation-responsive CREs. **a**, Enrichment of heritability in stimulation-responsive CREs in various cell-types. *Solid circles*, significant enrichments with FDR <0.05. **b**, Ranking of cell-type relevance to diseases based on heritability enrichment. Regression coefficient, from the analysis in **(a)**, can be interpreted as the extent of stimulation-specific heritability enrichment in a particular cell-type, and thus cell-type relevance. *Error bars*, standard error of the estimate. *Blue line* and *grey shade*, linear regression mean and 95% confidence intervals.

**Supplementary Fig. 13.**
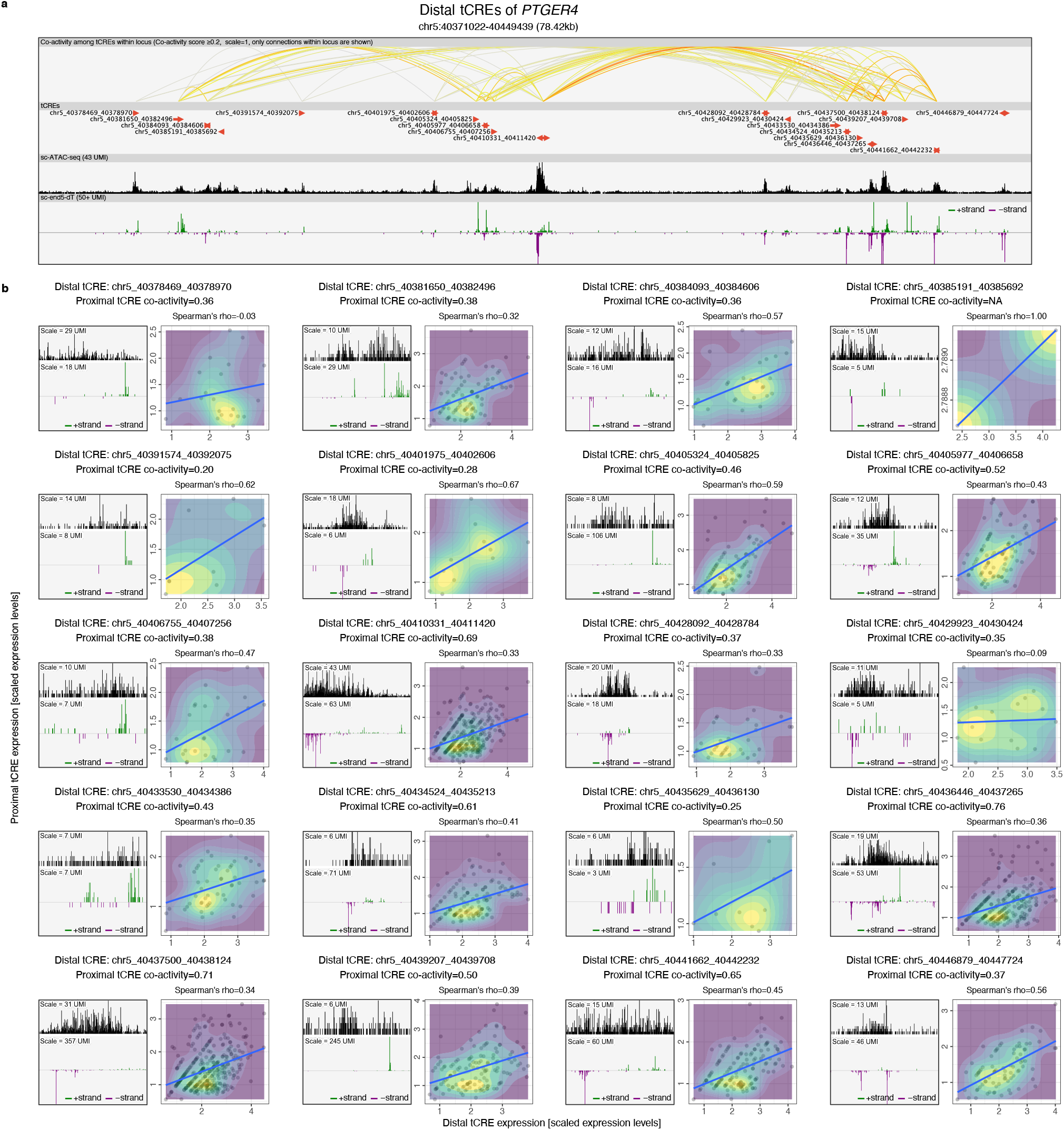
Distal tCRE activity at the *PTGER4* locus. **a**, Overview of the distal tCREs in close proximity to *PTGER4*. Twenty distal tCREs were shown. Co-activity among these 20 tCREs, with co-activity score ≥0.2, is represented by the color of the arcs. Only co-activity among tCRE within the view was shown. Resting and stimulated PBMC data were pooled in the sc-ATAC-seq and sc-end5-dT tracks. *Green* and *blue bars* in the sc-end5-dT track represent the + and – strand signal. The view was generated in the Zenbu genome browser with modifications (Data availability). **b**, Individual distal tCREs and their coactivity with *PTGER4* proximal tCRE. For each distal tCRE in **(a)**, a zoom-in view at the locus is shown. The scale of the signal bars is indicated as UMI counts. Expression of individual distal tCRE and the *PTGER4* proximal tCRE within single cells are plotted. Only cells with non-zero values in both tCREs are plotted. *Blue line*, mean of linear regression.

## References

1. Forrest, A. R. R. et al. A promoter-level mammalian expression atlas. Nature 507, 462–470 (2014).

2. Andersson, R. et al. An atlas of active enhancers across human cell types and tissues. Nature 507, 455–461 (2014).

3. Maurano, M. T. et al. Systematic Localization of Common Disease-Associated Variation in Regulatory DNA. Science 337, 1190–1195 (2012).

4. Heinz, S., Romanoski, C. E., Benner, C. & Glass, C. K. The selection and function of cell type-specific enhancers. Nat. Rev. Mol. Cell Biol. 16, 144–154 (2015).

5. Catarino, R. R. & Stark, A. Assessing sufficiency and necessity of enhancer activities for gene expression and the mechanisms of transcription activation. Genes Dev. 32, 202–223 (2018).

6. Boix, C. A., James, B. T., Park, Y. P., Meuleman, W. & Kellis, M. Regulatory genomic circuitry of human disease loci by integrative epigenomics. Nature 590, 300–307 (2021).

7. Trapnell, C. Defining cell types and states with single-cell genomics. Genome Res. 25, 1491–1498 (2015).

8. Wagner, A., Regev, A. & Yosef, N. Revealing the vectors of cellular identity with single-cell genomics. Nat. Biotechnol. 34, 1145–1160 (2016).

9. Buenrostro, J. D. et al. Single-cell chromatin accessibility reveals principles of regulatory variation. Nature 523, 486–490 (2015).

10. Cusanovich, D. A. et al. A Single-Cell Atlas of In Vivo Mammalian Chromatin Accessibility. Cell 174, 1309-1324.e18 (2018).

11. Chen, H. et al. Assessment of computational methods for the analysis of single-cell ATAC-seq data. Genome Biol. 20, 241 (2019).

12. Chen, S., Lake, B. B. & Zhang, K. High-throughput sequencing of the transcriptome and chromatin accessibility in the same cell. Nat. Biotechnol. 37, 1452–1457 (2019).

13. Cao, J. et al. Joint profiling of chromatin accessibility and gene expression in thousands of single cells. Science 361, 1380–1385 (2018).

14. Ma, S. et al. Chromatin Potential Identified by Shared Single-Cell Profiling of RNA and Chromatin. Cell 183, 1103-1116.e20 (2020).

15. Granja, J. M. et al. Single-cell multiomic analysis identifies regulatory programs in mixed-phenotype acute leukemia. Nat. Biotechnol. 37, 1458–1465 (2019).

16. Pliner, H. A. et al. Cicero Predicts cis-Regulatory DNA Interactions from Single-Cell Chromatin Accessibility Data. Mol. Cell 71, 858-871.e8 (2018).

17. Xiong, L. et al. SCALE method for single-cell ATAC-seq analysis via latent feature extraction. Nat. Commun. 10, 4576 (2019).

18. Thibodeau, A., Uyar, A., Khetan, S., Stitzel, M. L. & Ucar, D. A neural network based model effectively predicts enhancers from clinical ATAC-seq samples. Sci. Rep. 8, 16048 (2018).

19. Kim, T. H. et al. Analysis of the vertebrate insulator protein CTCF-binding sites in the human genome. Cell 128, 1231–1245 (2007).

20. Pang, B. & Snyder, M. P. Systematic identification of silencers in human cells. Nat. Genet. 52, 254–263 (2020).

21. Shiraki, T. et al. Cap analysis gene expression for high-throughput analysis of transcriptional starting point and identification of promoter usage. Proc. Natl. Acad. Sci. U. S. A. 100, 15776–15781 (2003).

22. Whalen, S., Truty, R. M. & Pollard, K. S. Enhancer-promoter interactions are encoded by complex genomic signatures on looping chromatin. Nat. Genet. 48, 488–496 (2016).

23. Rennie, S., Dalby, M., van Duin, L. & Andersson, R. Transcriptional decomposition reveals active chromatin architectures and cell specific regulatory interactions. Nat. Commun. 9, 487 (2018).

24. Mikhaylichenko, O. et al. The degree of enhancer or promoter activity is reflected by the levels and directionality of eRNA transcription. Genes Dev. 32, 42–57 (2018).

25. Kouno, T. et al. C1 CAGE detects transcription start sites and enhancer activity at single-cell resolution. Nat. Commun. 10, 360 (2019).

26. Cumbie, J. S., Ivanchenko, M. G. & Megraw, M. NanoCAGE-XL and CapFilter: an approach to genome wide identification of high confidence transcription start sites. BMC Genomics 16, 597 (2015).

27. Adiconis, X. et al. Comprehensive comparative analysis of 5’-end RNA-sequencing methods. Nat. Methods 15, 505–511 (2018).

28. Cheng, J. et al. Transcriptional Maps of 10 Human Chromosomes at 5-Nucleotide Resolution. Science 308, 1149–1154 (2005).

29. Kodzius, R. et al. CAGE: cap analysis of gene expression. Nat. Methods 3, 211–222 (2006).

30. Yang, L., Duff, M. O., Graveley, B. R., Carmichael, G. G. & Chen, L.-L. Genomewide characterization of non-polyadenylated RNAs. Genome Biol. 12, R16 (2011).

31. Hirabayashi, S. et al. NET-CAGE characterizes the dynamics and topology of human transcribed cis-regulatory elements. Nat. Genet. 51, 1369–1379 (2019).

32. La Manno, G. et al. RNA velocity of single cells. Nature 560, 494–498 (2018).

33. Gaidatzis, D., Burger, L., Florescu, M. & Stadler, M. B. Analysis of intronic and exonic reads in RNA-seq data characterizes transcriptional and post-transcriptional regulation. Nat. Biotechnol. 33, 722–729 (2015).

34. Cvetesic, N. et al. SLIC-CAGE: high-resolution transcription start site mapping using nanogram-levels of total RNA. Genome Res. 28, 1943–1956 (2018).

35. Kanamori-Katayama, M. et al. Unamplified cap analysis of gene expression on a single-molecule sequencer. Genome Res. 21, 1150–1159 (2011).

36. Affymetrix ENCODE Transcriptome Project & Cold Spring Harbor Laboratory ENCODE Transcriptome Project. Post-transcriptional processing generates a diversity of 5’-modified long and short RNAs. Nature 457, 1028–1032 (2009).

37. Cheng, J. et al. A role for H3K4 monomethylation in gene repression and partitioning of chromatin readers. Mol. Cell 53, 979–992 (2014).

38. Bae, S. & Lesch, B. J. H3K4me1 Distribution Predicts Transcription State and Poising at Promoters. Front. Cell Dev. Biol. 8, 289 (2020).

39. Preker, P. et al. PROMoter uPstream Transcripts share characteristics with mRNAs and are produced upstream of all three major types of mammalian promoters. Nucleic Acids Res. 39, 7179–7193 (2011).

40. Yan, F., Powell, D. R., Curtis, D. J. & Wong, N. C. From reads to insight: a hitchhiker’s guide to ATAC-seq data analysis. Genome Biol. 21, 22 (2020).

41. Stuart, T. et al. Comprehensive Integration of Single-Cell Data. Cell 177, 1888-1902.e21 (2019).

42. Schep, A. N., Wu, B., Buenrostro, J. D. & Greenleaf, W. J. chromVAR: inferring transcription-factor-associated accessibility from single-cell epigenomic data. Nat. Methods 14, 975–978 (2017).

43. Javierre, B. M. et al. Lineage-Specific Genome Architecture Links Enhancers and Non-coding Disease Variants to Target Gene Promoters. Cell 167, 1369-1384.e19 (2016).

44. Zhang, P. et al. Relatively frequent switching of transcription start sites during cerebellar development. BMC Genomics 18, 461 (2017).

45. Finucane, H. K. et al. Partitioning heritability by functional annotation using genome-wide association summary statistics. Nat. Genet. 47, 1228–1235 (2015).

46. Ni, G., Moser, G., Schizophrenia Working Group of the Psychiatric Genomics Consortium, Wray, N. R. & Lee, S. H. Estimation of Genetic Correlation via Linkage Disequilibrium Score Regression and Genomic Restricted Maximum Likelihood. Am. J. Hum. Genet. 102, 1185–1194 (2018).

47. Ramilowski, J. A. et al. Functional annotation of human long noncoding RNAs via molecular phenotyping. Genome Res. 30, 1060–1072 (2020).

48. Trynka, G. et al. Chromatin marks identify critical cell types for fine mapping complex trait variants. Nat. Genet. 45, 124–130 (2013).

49. Akdis, M. et al. T helper (Th) 2 predominance in atopic diseases is due to preferential apoptosis of circulating memory/effector Th1 cells. FASEB J. Off. Publ. Fed. Am. Soc. Exp. Biol. 17, 1026–1035 (2003).

50. Skapenko, A., Leipe, J., Lipsky, P. E. & Schulze-Koops, H. The role of the T cell in autoimmune inflammation. Arthritis Res. Ther. 7, S4 (2005).

51. Libioulle, C. et al. Novel Crohn disease locus identified by genome-wide association maps to a gene desert on 5p13.1 and modulates expression of PTGER4. PLoS Genet. 3, e58 (2007).

52. Grindberg, R. V. et al. RNA-sequencing from single nuclei. Proc. Natl. Acad. Sci. 110, 19802–19807 (2013).

53. Mercer, T. R. et al. Targeted RNA sequencing reveals the deep complexity of the human transcriptome. Nat. Biotechnol. 30, 99–104 (2012).

54. Severin, J. et al. Interactive visualization and analysis of large-scale sequencing datasets using ZENBU. Nat. Biotechnol. 32, 217–219 (2014).

55. Frankish, A. et al. GENCODE reference annotation for the human and mouse genomes. Nucleic Acids Res. 47, D766–D773 (2019).

56. Fort, A. et al. Deep transcriptome profiling of mammalian stem cells supports a regulatory role for retrotransposons in pluripotency maintenance. Nat. Genet. 46, 558–566 (2014).

57. Conrad, T. & Ørom, U. A. Cellular Fractionation and Isolation of Chromatin-Associated RNA. Methods Mol. Biol. Clifton NJ 1468, 1–9 (2017).

58. Murata, M. et al. Detecting expressed genes using CAGE. Methods Mol. Biol. Clifton NJ 1164, 67–85 (2014).

59. Buenrostro, J. D., Wu, B., Chang, H. Y. & Greenleaf, W. J. ATAC-seq: A Method for Assaying Chromatin Accessibility Genome-Wide. Curr. Protoc. Mol. Biol. 109, 21.29.1-21.29.9 (2015).

60. Kim, D., Paggi, J. M., Park, C., Bennett, C. & Salzberg, S. L. Graph-based genome alignment and genotyping with HISAT2 and HISAT-genotype. Nat. Biotechnol. 37, 907–915 (2019).

61. Korotkevich, G. et al. Fast gene set enrichment analysis. bioRxiv 060012 (2021) doi:10.1101/060012.

62. Butler, A., Hoffman, P., Smibert, P., Papalexi, E. & Satija, R. Integrating single-cell transcriptomic data across different conditions, technologies, and species. Nat. Biotechnol. 36, 411–420 (2018).

63. Hou, R., Denisenko, E. & Forrest, A. R. R. scMatch: a single-cell gene expression profile annotation tool using reference datasets. Bioinformatics 35, 4688–4695 (2019).

64. Fang, R. et al. SnapATAC: A Comprehensive Analysis Package for Single Cell ATAC-seq. bioRxiv 615179 (2020) doi:10.1101/615179.

65. Korsunsky, I. et al. Fast, sensitive and accurate integration of single-cell data with Harmony. Nat. Methods 16, 1289–1296 (2019).

66. Khan, A. et al. JASPAR 2018: update of the open-access database of transcription factor binding profiles and its web framework. Nucleic Acids Res. 46, D260–D266 (2018).

67. Kawaji, H. et al. Comparison of CAGE and RNA-seq transcriptome profiling using clonally amplified and single-molecule next-generation sequencing. Genome Res. 24, 708–717 (2014).

68. Li, H. et al. The Sequence Alignment/Map format and SAMtools. Bioinforma. Oxf. Engl. 25, 2078–2079 (2009).

69. Frith, M. C. et al. A code for transcription initiation in mammalian genomes. Genome Res. 18, 1–12 (2008).

70. Kuhn, M. Building Predictive Models in R Using the caret Package. J. Stat. Softw. 28, (2008).

71. Quinlan, A. R. & Hall, I. M. BEDTools: a flexible suite of utilities for comparing genomic features. Bioinforma. Oxf. Engl. 26, 841–842 (2010).

72. Bulik-Sullivan, B. K. et al. LD Score regression distinguishes confounding from polygenicity in genome-wide association studies. Nat. Genet. 47, 291–295 (2015).

73. Li, M. J. et al. GWASdb v2: an update database for human genetic variants identified by genome-wide association studies. Nucleic Acids Res. 44, D869–876 (2016).

74. Buniello, A. et al. The NHGRI-EBI GWAS Catalog of published genome-wide association studies, targeted arrays and summary statistics 2019. Nucleic Acids Res. 47, D1005–D1012 (2019).

75. Purcell, S. et al. PLINK: a tool set for whole-genome association and population-based linkage analyses. Am. J. Hum. Genet. 81, 559–575 (2007).

76. de Leeuw, C. A., Mooij, J. M., Heskes, T. & Posthuma, D. MAGMA: generalized gene-set analysis of GWAS data. PLoS Comput. Biol. 11, e1004219 (2015).

77. Finucane, H. K. et al. Heritability enrichment of specifically expressed genes identifies disease-relevant tissues and cell types. Nat. Genet. 50, 621–629 (2018).

78. Wickham, H. ggplot2: Elegant Graphics for Data Analysis. (Springer-Verlag New York, 2016).

